# *Arabidopsis thaliana* ACTIN DEPOLYMERIZING FACTORs are novel susceptibility factors for *Colletotrichum higginsianum*

**DOI:** 10.64898/2026.09.01.748704

**Authors:** Miku Ohashi, Sakura Aoki, Takashi L. Shimada, Takashi Ueda, Masaaki Umeda, Noriko Inada

**Affiliations:** Graduate School of Agriculture, Osaka Metropolitan University, 1-1 Gakuen-cho, Naka-ku, Sakai, Osaka 5998531, Japan; Graduate School of Life and Environmental Sciences, Osaka Prefecture University, 1-1 Gakuen-cho, Naka-ku, Sakai, Osaka 5998531, Japan; Graduate School of Horticulture, Chiba University, 648 Matsudo, Matsudo, Chiba 271-8510, Japan; Research Center for Space Agriculture and Horticulture, Chiba University, 648 Matsudo, Matsudo, Chiba 271-8510, Japan; Division of Cellular Dynamics, National Institute for Basic Biology, Nishigonaka 38, Myodaijim Okazaki, Aichi 444-8585, Japan; Basic Biology Program, Graduate Institute for Advanced Studies, SOKENDAI, Nishigonaka 38, Myodaiji, Okazaki, Aichi 444-8585, Japan; Graduate School of Science and Technology, Nara Institute of Science and Technology, Takayama, Ikoma, Nara 630-0001, Japan

**Keywords:** ACTIN DEPOLYMERIZING FACTOR, actin filaments, *Arabidopsis thaliana*, *Colletotrichum higginsianum*, PEN2

## Abstract

*Colletotrichum higginsianum* (*Ch*) is a hemibiotrophic fungal pathogen that infects Brassicaceae plants, including *Arabidopsis thaliana*. The molecular mechanisms underlying the *Ch*-*A. thaliana* interaction are not fully understood. Particularly, the susceptibility factor against *Ch* infection remains to be determined. Here, we report that *A. thaliana* ACTIN DEPOLYMERIZING FACTORs (ADFs), ancient proteins that regulate the organization and dynamics of actin filaments (AFs), function as susceptibility factors during *Ch* infection. Among 11 *ADF*s encoded in *A. thaliana* genome, subclass I *ADF*s that include *ADF1*, *-2*, *-3*, and *-4*, express throughout the plant. We found that knockout mutant of *ADF4* and transgenic plants in which the expression of all of subclass I members is suppressed (*ADF1-4Ri*) exhibited increased resistance to *Ch*. Cytological analyses revealed that both *Ch* penetration and secondary hyphae formation were suppressed in *adf4* and *ADF1-4Ri*. This enhanced resistance was associated with suppression of *Ch*-induced AF fragmentation. In addition, we found that PENETRATION 2 (PEN2) plays a critical role in the *Ch* resistance in *adf4* and *ADF1-4Ri*. Our findings suggest that subclass I ADFs promote AF fragmentation during *Ch* infection, thereby suppressing PEN2-associated mitochondria accumulation at *Ch* entry sites. Together, these results raise the possibility that *Ch* exploits host ADF-dependent actin regulation to facilitate successful infection.

## INTRODUCTION

*Colletotrichum* species are among the most important fungal pathogens causing anthracnose disease in a wide range of crops, fruits, and ornamental plants (Dean *et al*., 2012). *C. higginsianum* (*Ch*) infects various Brassicaceae plants, including *B. napus*, *B. oleracea* (cabbage), *B. campestris* (turnip and pak-choi), *Raphanus sativus* (radish), and *Sinapis alba* (white mustard), as well as members of the Fabaceae, including *Lentis culinalis* and *Vigna unguiculata* (O’Connell *et al*., 2004). *Ch* also infects the model plant *Arabidopsis thaliana*, and the *Ch*-*A. thaliana* interaction has been widely used as a model pathosystem to investigate the molecular mechanisms by which fungal pathogens establish infection in host plants and how host plants respond to infection (Yan *et al*., 2018).

*Ch* has a hemibiotrophic lifestyle and establishes infection in host *A. thaliana* through the development of a series of specialized infection structures. After *Ch* conidia are inoculated onto *A. thaliana* leaves, the germinated conidia develop appressoria within 24 hours post inoculation (hpi). These darkly pigmented appressoria accumulate osmolytes and thereby generate high turgor pressure, enabling mechanical penetration of the host cuticle and cell wall. Following penetration, an infection peg emerging from the appressorium expands to form biotrophic bulbous primary hyphae in the apoplast. This biotrophic stage is restricted to the initially invaded cells. The fungus subsequently switches to a necrotrophic phase, which is characterized by the formation of thin secondary hyphae. These necrotrophic hyphae invade adjacent cells and colonize the surrounding tissue. At 4∼6 days post inoculation (dpi), necrotic lesions become readily visible at the inoculation sites (Narusaka *et al*., 2004; O’Connell *et al*., 2004).

To establish successful infection, pathogens must evade multiple layers of host immunity. The first layer of plant immunity is triggered by recognition of pathogen-associated molecular patterns (PAMPs). Plants can also recognize endogenous signals generated by cellular damage, known as damage-associated molecular patterns (DAMPs) (Saijo *et al*., 2018). Immunity activated by the recognition of PAMPs or DAMPs is referred to as PAMP-triggered immunity (PTI). A second layer of immunity is activated through recognition of pathogen effectors and is therefore termed effector-triggered immunity (ETI) (Jones & Dangl 2006). Extensive manipulation of host cellular functions by pathogen effectors is thought to occur during interactions between living host plants and biotrophic pathogens. A previous analysis of the *Ch* genome identified 365 candidate secreted effectors (CSEPs) predicted to function extracellularly and to be unique to the *Colletotrichum* genus (O’Connell *et al*., 2012). Among these CSEPs, ChELP1 and ChELP2 function during the biotrophic stage of infection. These proteins contain LysM domains that bind fungal chitin or peptidoglycans, thereby suppressing chitin-triggered PTI (Takahara *et al*., 2016). Screening of T-DNA insertion mutants has also identified several *Ch* pathogenicity factors, including importin β2, ATP-binding endoribonuclease, β-1,3(4)-glucanase, and plasma membrane H+-ATPase (summarized in Yan *et al*., 2018). Understanding host susceptibility factors is equally important for elucidating host-pathogen interactions. However, little is currently known about host susceptibility factors involved in the *Ch*-*A. thaliana* interaction.

In this study, we identified *A. thaliana* ACTIN DEPOLYMERIZING FACTORs (ADFs) as novel regulators of susceptibility to *Ch*. ADF is an evolutionarily conserved protein found throughout eukaryotes that functions in the depolymerization and severing of actin microfilaments (AFs) (Inada, 2017). ADFs have been reported to regulate host responses to a variety of pathogens (Sun *et al*., 2023). *A. thaliana ADF4* functions in resistance to the bacterial pathogen *Pseudomonas syringae* pv. *tomato* (*Pst*) DC3000 carrying AvrPphB. Loss of *ADF4*, but not *ADF1* or *ADF3*, increases susceptibility to *Pst* DC3000 AvrPphB (Tian *et al*., 2009). Expression of *A. thaliana ADF2* is upregulated in galls induced by the root-knot nematode *Meloidogyne incognita*, and suppression of *ADF2* expression arrests gall development (Clément *et al*., 2009). Loss of *A. thaliana ADF3* increases susceptibility to the green peach aphid *Myzus persicae* (Mondal *et al*., 2018). Wheat *TaADF4* and *TaADF7* function in resistance to avirulent *Puccinia striiformis* f. sp. *tritici* pathotypes. Suppression of *TaADF4* (Zhang *et al*., 2017) or *TaADF7* (Fu et al., 2014) increases susceptibility to avirulent *P. striiformis*. In contrast, suppression of *TaADF3* enhances resistance to both virulent and avirulent *P. striiformis* pathotypes and therefore functions in susceptibility to *P. striiformis* (Tang *et al*., 2016). Similarly, suppression of cotton (*Gossypium hirsutum*) *GhADF6* expression increases resistance to the vascular fungal pathogen *Verticillium dahlia* (Sun *et al*., 2021b).

Land plants have evolved a family of ADFs, and the *A. thaliana* genome contains 11 ADF members that are classified into four subclasses. Subclass I ADFs, including *ADF1*, *ADF2*, *ADF3*, and *ADF4*, are expressed throughout the plant at relatively high levels (Ruzicka *et al*., 2007). We previously showed that subclass I ADFs, particularly *ADF4*, function in the response to the *A. thaliana*-adapted powdery mildew fungus *Golovinomyces orontii* (*Go*). At 2 weeks post inoculation, wild-type Col-0 leaves are almost completely covered with whitish *Go* mycelia, whereas mycelial development is significantly suppressed in *adf4*. This resistance is further enhanced in transgenic plants in which expression of all four subclass I ADF genes is suppressed (*ADF1-4Ri*; Tian *et al*., 2009; Inada *et al*., 2016). Although the primary function of ADF is the regulation of AFs at the cell surface, neither *adf4* nor *ADF1-4Ri* exhibited significant alterations in AF organization relative to wild type. ADF4 was found to localize both to cytoplasmic AFs and to the nucleus, and complementation assays using ADF4 fused to a nuclear export signal (NES) demonstrated that nuclear localization of ADF4 is important for the response to *Go* (Inada *et al*., 2016).

In this study, we found that *adf4* and *ADF1-4Ri* also exhibit increased resistance to *Ch*. This enhanced resistance was evident during both the early stage of infection, when *Ch* penetrates host epidermal cells, and the later stage, when secondary hyphae are produced. Several lines of evidence indicated that the mechanism underlying increased *Ch* resistance in *adf4* and *ADF1-4Ri* differs from that responsible for increased *Go* resistance in these plants. A recent study reported that *Ch* infection induces fragmentation of AFs in infected *A. thaliana* cells (Shimada *et al*., 2026). Therefore, we performed a detailed analysis of AF organization in *Ch*-infected cells. We also examined the function of PEN2, which plays a role in penetration resistance against non-adapted *Colletotrichum* species (Hiruma *et al*., 2010). Our results strongly suggest that *Ch* manipulates ADF function to facilitate successful infection.

## MATERIALS AND METHODS

### Plant materials and growth condition

*adf3* (SALK_139265), *adf4* (Garlic_823_A11b.1b.Lb3Fa) and *ADF1-4Ri* lines were described previously (Tian *et al*., 2009). *act2-1/act8-2* double mutant and *act7-4* single mutant were described previously (Kandasamy *et al*., 2009). *pen2-1* was described previously (Collins *et al*., 2003; Lipka *et al*., 2005; Irieda & Takano, 2021). *A. thaliana* lines expressing GFP-ABD2 (Shimono *et al*., 2016), PEN2-GFP (Lipka *et al*., 2005), GFP-PTS1 (Mano *et al*., 2002) were described previously.

*A. thaliana* seeds were suspended in autoclaved 0.1% agarose and incubated at 4°C for vernalization for more than 1 day (up to 2 weeks) before direct sowing on 1:3 metromix:vermiculite in plastic pots. Entire pots were covered with plastic wrap for 1 week after sowing to maintain humidity and to encourage germination. Plants were grown at 22°C in a growth chamber under a 12-h light:12-h dark photoperiod (LH-411PFD-S, NK Systems, Osaka, Japan).

### Fungal strain and growth

*Colletotrichum higginsianum* isolate (MAFF305635) was obtained from MAFF Genbank, Japan. Cultures of the isolate was maintained on 3.9% (w/v) potato dextrose agar medium (Shimadzu Diagnostics Corporation, Tokyo, Japan) at 24°C in the dark.

### Fungal inoculation

For macroscopic and microscopic observation, mature leaves of 4 week old *A. thaliana* plants or cotyledons of 2 week old plants were inoculated with 5 µL drops of *Ch* conidial suspension that contained 2.5 x 10^5^ conidia per mL. Inoculated plants were incubated in a growth chamber at 22°C with a 12-h light:12-h dark photoperiod at 100% humidity (LH-411PFD-S, NK Systems, Osaka, Japan).

### Macroscopic observation of *Ch* lesions

Leaves at 5-6 days post inoculation was excised and observed with a stereo microscope (Leica MZ10F, Leica Microsystems, Germany) equipped with CCD camera (Olympus DP73, Olympus, Tokyo, Japan) and an image capture software (CellSens Standard, Olympus, Japan). The size of each lesion was determined by analyzing the obtained images with ImageJ.

### RT-qPCR

*Ch*-infected mature leaves were harvested and frozen in liquid N_2_. Total RNA was extracted using Sepasol-RNA I Super G (Nacalai tesque, Kyoto, Japan) in accordance with the manufacturer’s protocol. cDNA was synthesized with the ReverTra Ace qPCR RT Master Mix with gDNA Remover (TOYOBO, Osaka, Japan) using 100 ng of total RNA. RT-qPCR was carried out using THUNDERBIRD SYBR qPCR Mix (TOYOBO, Osaka, Japan) on 7300 Real Time PCR System (Applied Biosystems/Thermo Fisher Scientific, Japan). *UBQ11* (At4g05050) was used as internal control gene for normalization. Primer information is shown in Table S1.

### Histochemical staining and light microscopy

For observation of *Ch* invasion and formation of hyphae, *Ch*-inoculated leaves were immersed with lactophenol-trypan blue solution that contained 10 mL of lactic acid, 10 mL of glycerol, 10 g of granular phenol, 10 mg of trypan blue and 10 mL of milliQ water. Leaves in the lactophenol-trypan blue solution were boiled for 1 min, then destained with saturated chloral hydrate (2.5 g/mL). For observation of ROS accumulation, leaves were stained with DAB solution as described previously (Inada *et al*., 2016). Stained leaves were examined with a light microscope (ECLIPSE E600, Nikon, Japan) equipped with 40x objective lens (Plant Fluor 40x N.A. 0.75, Nikon) and CCD camera (ORCA-ER, Hamamatsu photonics, Japan).

### Observation of intracellular structures with confocal laser scanning microscope

To observe AFs, we crossed GFP-ABD2 expressing Col-0 with *adf4* and *ADF1-4Ri*. Cotyledons of GFP-ABD2 expressing Col-0, *adf4* and *ADF1-4Ri* plants were inoculated with *Ch* and observed using confocal laser scanning microscope (CLSM, LSM700, Zeiss, Germany) equipped with 20x objective lens (Plan-Apochromat 20x DICII N.A. 0.8, Zeiss, Germany). GFP was excited with a 488 nm argon laser, and the emitted fluorescence was filtered with 505-600 nm bandpass filter.

For observation of mitochondria, *Ch*-infected leaves were immersed with MitoTracker Green FM (M7514, Thermo Fisher Scientific, Japan) that was diluted to 500 nM with 1/2 MS medium, vacuum-infiltrated for 5 min, and incubated for 15 min in the dark. The fluorescence images of MitoTracker Green FM were obtained using CLSM (LSM700, Zeiss, Germany) equipped with 20x objective lens (Plan-Apochromat 20x DICII N.A. 0.8, Zeiss, Germany). GFP was excited with a 488 nm argon laser, and the emitted fluorescence was filtered with 505-600 nm bandpass filter.

## RESULTS

### Resistance against *Ch* was increased in *adf4* and *ADF1-4Ri*

We previously showed that *adf4* exhibits increased resistance to the obligate biotrophic fungus *Go*, and that the level of *Go* resistance is even greater in *ADF1-4Ri* than in *adf4* (Inada *et al*., 2016). In *adf4*, and more prominently in *ADF1-4Ri*, reactive oxygen species (ROS) accumulate in Go-infected cells. This ROS accumulation likely induces cell death in infected cells, thereby suppressing fungal nutrient acquisition through the specialized feeding structure known as a haustorium, and consequently inhibiting hyphal development and conidiophore formation (Inada *et al*., 2016). Because *Ch* undergoes a biotrophic phase during the early stage of infection, we hypothesized that *adf4* and *ADF1-4Ri* might also exhibit enhanced resistance to *Ch*.

To test this hypothesis, we inoculated Col-0, *adf4*, and *ADF1-4Ri* with *Ch* conidia and examined lesion development at 6 days post inoculation (dpi). As shown in Fig. 1a, the *A. thaliana*-adapted *Ch* isolate formed necrotic lesions on Col-0 leaves at 6 dpi. In contrast, lesion size was significantly reduced in both *adf4* and *ADF1-4Ri* (Fig. 1a). Quantification revealed reductions of 57% in *adf4* and 60% in *ADF1-4Ri* relative to wild type (Fig. 1b). The enhanced resistance of *adf4* and *ADF1-4Ri* was further confirmed by RT-qPCR analysis comparing the expression level of *ChACT* with that of the host *A. thaliana UBQ11* gene in *Ch*-infected leaves at 6 dpi (Fig. 1c). Expression of *ChACT* was reduced by 90% in *adf4*, and almost no amplification was detected in *ADF1-4Ri*, indicating that *Ch* growth was strongly suppressed in these plants.

**Figure 1.**
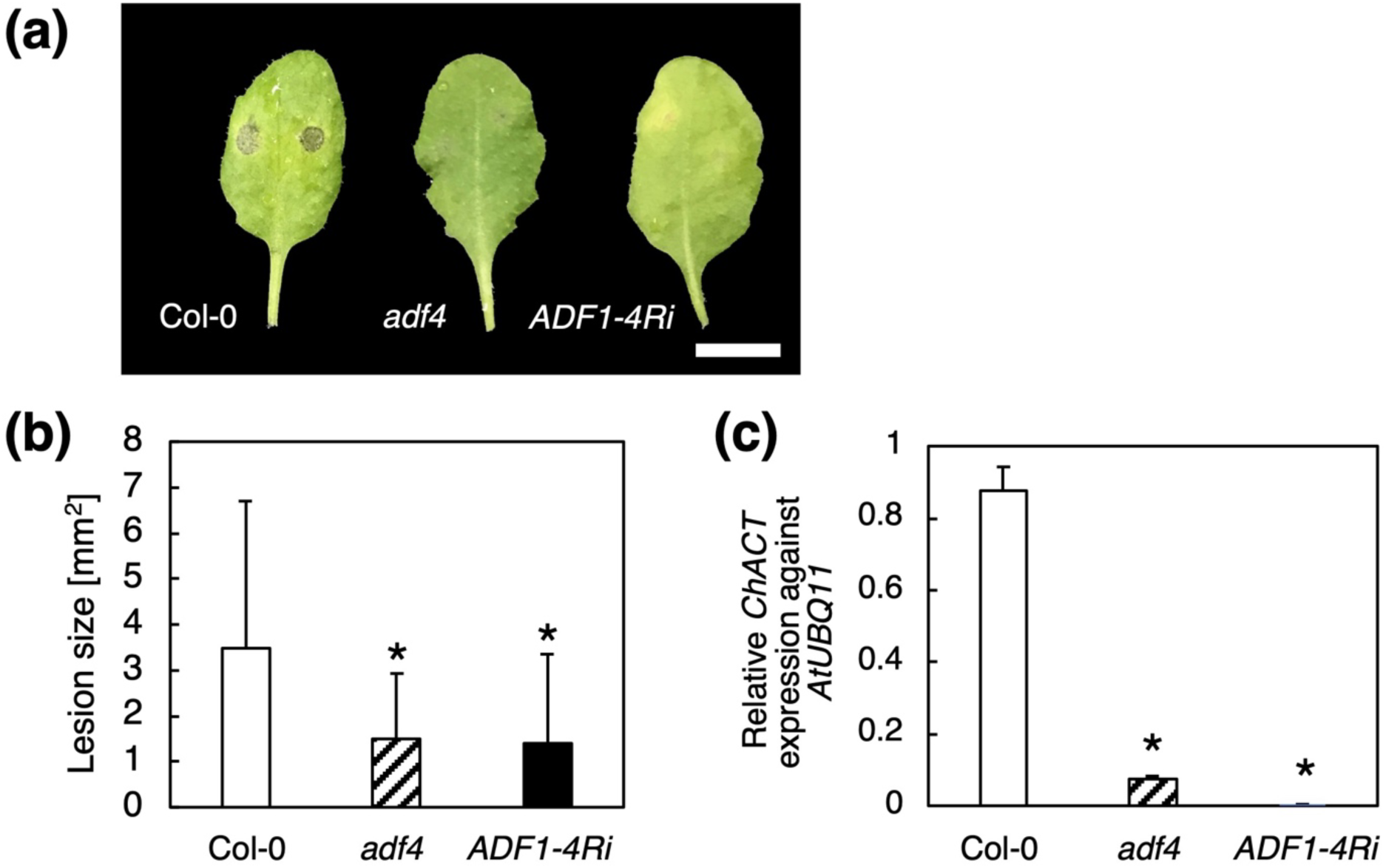
*Arabidopsis thaliana* subclass I ADFs are involved in the response against adapted *Colletotrichum higginsianum* (*Ch*). (**a**) Col-0, *adf4*, and *ADF1-4Ri* mature leaves inoculated with *Ch* at 6 days post inoculation (dpi). Bar indicates 5 mm. (**b**) The size of *Ch*-induced lesion at 5 dpi. The data showed average and standard deviation of 29 lesions for Col-0, *adf4*, and *ADF1-4Ri* each. (**c**) Quantification of *Ch* in planta by qRT-PCR. For (**b**) and (**c**), experiments were repeated three times with similar results. The representative results are shown. Comparison with Col-0 was performed by Student’s *t*-test. Asterisks indicate that there is a significant difference (\**P*<0.05).

Among the four subclass I ADF members in *A. thaliana*, *ADF3* is expressed at the highest level (Ruzicka *et al*., 2007). We previously reported that an *adf3* knockout mutant also exhibited increased resistance to *Go*, although the resistance was weaker than that observed in *adf4*. While *adf3* displayed enhanced leaf yellowing following *Go* infection, a phenotype associated with increased resistance, *Go* mycelial development at 2 weeks post inoculation remained comparable to that in Col-0 (Inada *et al*., 2016). We found that the *adf3* knockout mutant also exhibited reduced *Ch* lesion size (Fig. S1). However, the increase in resistance was again less pronounced than that observed in *adf4*; *Ch* infection was reduced by 30% based on lesion size measurements and by 53% based on RT-qPCR analysis.

In *ADF1-4Ri* plants, expression of all subclass I ADF members is suppressed by RNAi (Tian *et al*., 2009). Four independent *ADF1-4Ri* lines, #1-4, #2-1, #3-2, and #4-2, have been generated, and line #2-1 was used for the experiments shown in Fig. 1. The degree of suppression of each subclass I ADF member differs among these lines (Tian *et al*., 2009). We confirmed that the other three lines also exhibited reduced lesion development at 6 dpi (Fig. S2). Furthermore, lesion size was reduced by 79%, 82%, and 74% in *ADF1-4Ri*#1-4, #3-2, and #4-2, respectively.

Taken together, suppression of subclass I *ADF* expression increased resistance to *Ch*. In this study, we used *adf4* and *ADF1-4Ri* to investigate the role of subclass I ADFs in the response to *Ch*. For *ADF1-4Ri*, line #2-1 was used for all subsequent analyses.

### Both *Ch* invasion and secondary hyphae formation were suppressed in *adf4* and *ADF1-4Ri*

Resistance to pathogens can be expressed at different stages of infection. To determine which stages of *Ch* infection were affected in *adf4* and *ADF1-4Ri*, we performed microscopic analyses. *Ch*-infected leaves were collected at 1, 2, and 3 dpi, fixed, stained with trypan blue, and observed microscopically (Fig. 2a-c). At 1 dpi, most *Ch* conidia had formed melanized appressoria (Fig. 2a). Following successful invasion, *Ch* produced biotrophic primary hyphae between the host cell wall and plasma membrane at 2-3 dpi (Fig. 2b). By 3 dpi, secondary hyphae had formed in a subset of conidia that had already developed primary hyphae (Fig. 2c).

**Figure 2.**
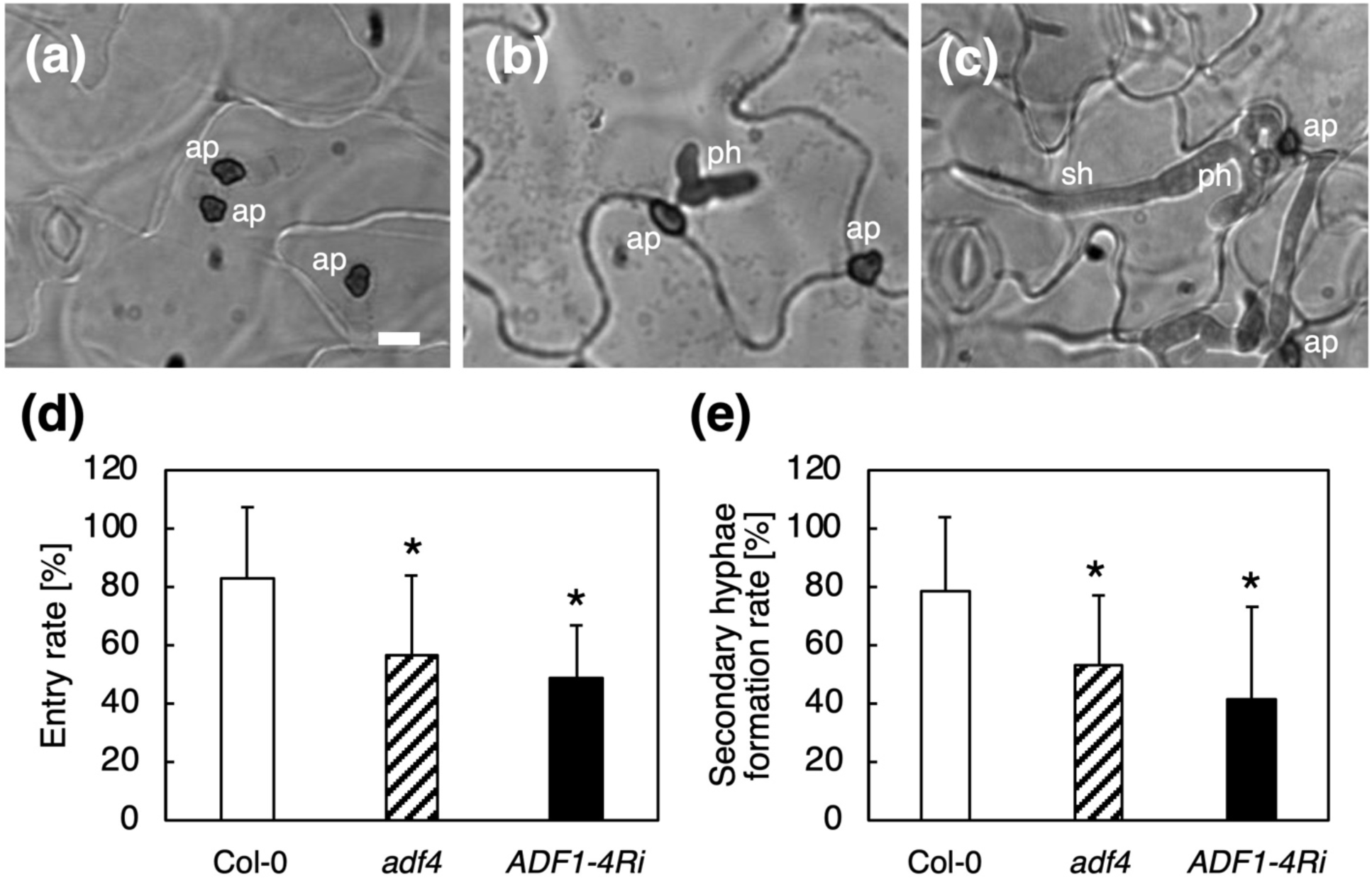
Both *Ch* entry and secondary hyphae formation were suppressed in *adf4* and *ADF1-4Ri*. (**a-c**) Col-0 mature leaves at 3 dpi were treated with lactophenol-trypan blue solution and observed under a microscope. Bar indicates 10 µm. ap, appressorium; ph, primary hyphae; sh, secondary hyphae. (**d**) *Ch* entry rate at 3 dpi. The percentage of appressoria with primary hyphae was counted for 20 appresoria per a leaf, and 18 leaves were used to calculate averaged percentages of entry. (**e**) The secondary hyphae formation rate at 3 dpi. The percentage of *Ch* forming secondary hyphae was determined among *Ch* forming primary hyphae. 20 *Ch* were counted per a leaf, and 18 leaves were used to calculate averaged percentages of secondary hyphae formation. For (**d**) and (**e**), comparison with Col-0 was performed by Student’s *t*-test. Asterisks indicate that there is a significant difference (* *P*<0.05).

We first examined the *Ch* entry rate in Col-0, *adf4*, and *ADF1-4Ri*. Entry rate was calculated as the percentage of conidia with primary hyphae among all conidia bearing appressoria. At 2 dpi, entry rates were 3.6% in Col-0, 1.1% in *adf4*, and 3.0% in *ADF1-4Ri*, with no statistically significant differences relative to Col-0 (Fig. S3). By 3 dpi, more than 80% of conidia with appressoria had successfully invaded host cells and formed primary hyphae in Col-0 (Fig. 2d). In contrast, entry rates were reduced to 57% in *adf4* and 49% in *ADF1-4Ri*. Relative to Col-0, this corresponded to reductions of 32% and 41%, respectively. These results indicate that resistance to *Ch* entry was significantly enhanced in both *adf4* and *ADF1-4Ri*.

During the analysis of *Ch* entry at 3 dpi, we also noticed a reduction in secondary hyphae formation in *adf4* and *ADF1-4Ri*. The frequency of secondary hyphae formation was determined by calculating the percentage of conidia with secondary hyphae among all conidia that had formed primary hyphae. In Col-0, nearly 80% of conidia with primary hyphae had developed secondary hyphae by 3 dpi (Fig. 2e). This percentage was significantly lower in both *adf4* and *ADF1-4Ri*, showing reductions of 32% and 47%, respectively, relative to Col-0 (Fig. 2e). Taken together, both resistance to *Ch* entry and suppression of secondary hyphae formation were enhanced in *adf4* and *ADF1-4Ri*.

### The mechanism underlying increased *Ch* resistance differed from that underlying increased *Go* resistance in *adf4* and *ADF1-4Ri*

We next examined whether the mechanism responsible for increased *Ch* resistance in *adf4* and *ADF1-4Ri* was the same as that underlying increased *Go* resistance in these plants. Previously, we showed that cell death associated with ROS accumulation in *Go*-infected cells was enhanced in *adf4* and *ADF1-4Ri* (Inada *et al*., 2016). To determine whether ROS accumulation during the biotrophic stage of *Ch* infection contributes to the increased *Ch* resistance observed in *adf4* and *ADF1-4Ri*, we performed DAB staining of *Ch*-infected leaves. Quantification of DAB-stained *Ch*-infected cells (Fig. S4a) in leaves collected at 3 dpi revealed that only 3% of cells containing *Ch* primary hyphae accumulated ROS in Col-0, and this percentage was similar in *adf4* and *ADF1-4Ri* (Fig. S4b). These results indicate that the increased *Ch* resistance in *adf4* and *ADF1-4Ri* is unlikely to result from host cell death-mediated suppression of biotrophic fungal activity.

In our previous study of the response to *Go*, we demonstrated that nuclear localization of ADF4 is important for resistance (Inada *et al*., 2016). Using GFP-hTalin-expressing plants (Takemoto et al., 2003), we examined the organization of actin microfilaments (AFs) at the cell surface and found that AF density and bundling in *adf4* and *ADF1-4Ri* were not significantly altered relative to Col-0 in uninfected cells. *ADF1-4Ri* exhibited significantly increased AF density at early time point (12-18 hpi) and decreased AF density at later time point (24-30 hpi), however, AF organization in *adf4* at those time points was comparable to that in Col-0. In addition, because ADF4-GFP localized to both the cytoplasm and the nucleus, we performed complementation analyses using ADF4-GFP and ADF4-GFP fused to a NES. While expression of ADF4-GFP complemented the increased *Go* resistance of *adf4*, *adf4* plants expressing ADF4-GFP-NES retained their enhanced resistance to *Go*. These findings demonstrated that the nuclear function of ADF4, rather than its role in regulating AFs at the cell surface, is important for the response to *Go* (Inada *et al*., 2016).

To determine whether the nuclear function of ADF4 also contributes to the response to *Ch*, we inoculated ADF4-GFP/*adf4* and ADF4-GFP-NES/*adf4* plants with *Ch* conidia and analyzed lesion development at 6 dpi together with *adf4* and Col-0 controls. Consistent with the results shown in Fig. 1, *adf4* exhibited a significant reduction in lesion size. In contrast, lesion sizes in both ADF4-GFP/*adf4* and ADF4-GFP-NES/*adf4* were comparable to those in Col-0 (Fig. S5). These results indicate that the nuclear function of ADF4 is not required for the response to *Ch*.

Taken together, these findings demonstrate that the mechanism underlying increased *Ch* resistance in *adf4* and *ADF1-4Ri* differs from that responsible for increased *Go* resistance.

### Fragmentation of AFs in *Ch*-infected cells was suppressed in *adf4* and ADF1-4Ri

A recent study showed that *Ch* infection induces fragmentation of AFs in infected cells (Shimada *et al*., 2026). Artificial induction of AF fragmentation by latrunculin B treatment increased the rate of *Ch* invasion and consequently enhanced susceptibility to *Ch*. Neither PAMP- nor DAMP-mediated signaling pathways were involved in the induction of AF fragmentation. This response was specifically triggered by *Ch* inoculation and was not observed following inoculation with the non-adapted species *C. tropicale*. *Ch*-induced AF fragmentation was observed in all tested host plants, including *B. oleracea* var. *capitata* (cabbage), *B. oleracea* var. *italica* (broccoli), and *B. rapa* var. *perviridis* (komatsuna). Interestingly, *Ch* also induced AF fragmentation in the *Ch*-resistant species *Cucumis sativus* (cucumber), whereas the cucumber-adapted pathogen *C. orbiculare* did not induce AF fragmentation in cucumber cells (Shimada *et al*., 2026).

To investigate whether loss of ADFs affects AF fragmentation during *Ch* infection, we used *A. thaliana* lines in which AFs are visualized with fluorescent protein. In our previous study of the *A. thaliana*-*Go* interaction, we employed GFP-hTalin-expressing plants (Takemoto *et al*., 2003) because GFP-hTalin was stably expressed in mature leaf epidermal cells, sites for *Go* infection. Furthermore, GFP-hTalin expression did not affect the increased *Go* resistance observed in *adf4* and *ADF1-4Ri* (Inada *et al*., 2016). In contrast, we found that GFP-hTalin expression substantially impaired the increased *Ch* resistance of *adf4* and *ADF1-4Ri*; lesion sizes at 6 dpi in GFP-hTalin/*adf4* and GFP-hTalin/*ADF1-4Ri* were comparable to those in GFP-hTalin/Col-0 (Fig. S6). Therefore, we chose the GFP-ABD2 line, which had previously been used to analyze AF fragmentation in *Ch*-infected cells (Shimada *et al*., 2026).

Plants expressing GFP-ABD2 displayed relatively strong GFP fluorescence in cotyledons, whereas fluorescence intensity decreased substantially during later developmental stages. Because *ADF4* expression is lower in cotyledons than in mature leaves (Ruzicka *et al*., 2007), and because we previously found that the altered nuclear morphology phenotype of *adf4* was less pronounced in cotyledons than in mature leaves (Matsumoto *et al*., 2023), we first examined whether the increased *Ch* resistance observed in mature leaves of *adf4* and *ADF1-4Ri* was retained in cotyledons. Both GFP-ABD2/*adf4* and GFP-ABD2/*ADF1-4Ri* exhibited significant reductions in *Ch* entry and secondary hyphae formation at 3 dpi relative to GFP-ABD2/Col-0 (Fig. S7). Thus, the increased *Ch* resistance of *adf4* and *ADF1-4Ri* was maintained in cotyledons. In addition, these results demonstrated that GFP-ABD2 expression did not interfere with the enhanced *Ch* resistance observed in *adf4* and *ADF1-4Ri*.

Cotyledons exhibiting strong GFP-ABD2 fluorescence were then subjected to *Ch* infection and examined using confocal laser scanning microscopy. Consistent with the previous report (Shimada *et al*., 2026), *Ch*-infected GFP-ABD2/Col-0 cells contained short fragmented AFs (Fig. 3a). The proportion of *Ch*-infected cells exhibiting fragmented AFs was 71% at 2 dpi and increased to 81% at 3 dpi in GFP-ABD2/Col-0 (Fig. 3b). In contrast, many *Ch*-infected cells in GFP-ABD2/*adf4* and GFP-ABD2/*ADF1-4Ri* retained bundled AFs and did not exhibit AF fragmentation (Fig. 3a). These bundled AFs did not show obvious polarization toward *Ch* entry sites. The percentage of *Ch*-infected cells with fragmented AFs was significantly lower in GFP-ABD2/*adf4* and GFP-ABD2/*ADF1-4Ri* than in GFP-ABD2/Col-0 at both 2 and 3 dpi (Fig. 3b). Relative to GFP-ABD2/Col-0, AF fragmentation was reduced by 40% in GFP-ABD2/*adf4* and 62% in GFP-ABD2/*ADF1-4Ri* at 2 dpi, and by 51% in GFP-ABD2/*adf4* and 78% in GFP-ABD2/*ADF1-4Ri* at 3 dpi. Overall, these results demonstrate that loss of ADF suppresses the AF fragmentation induced by *Ch* infection.

**Figure 3.**
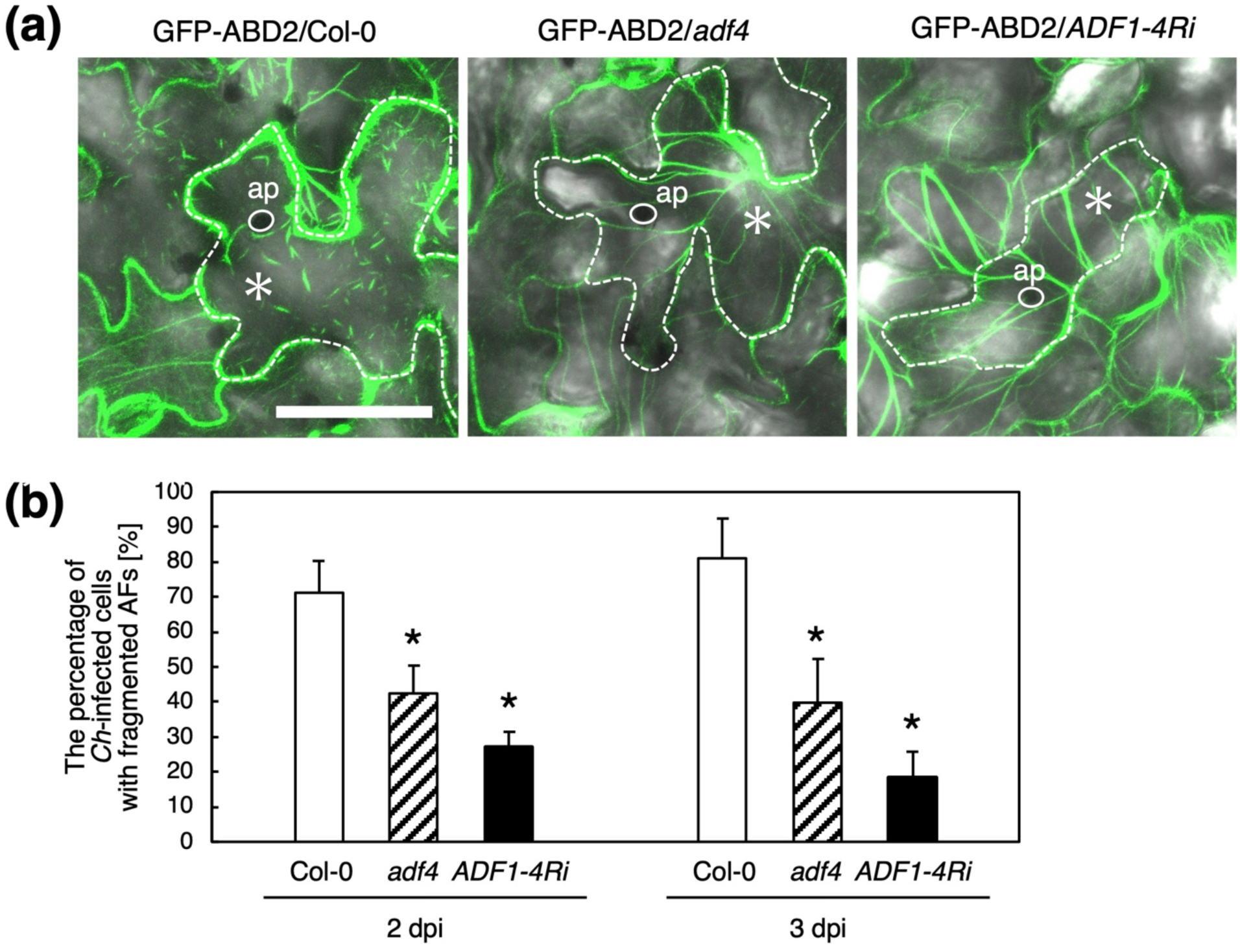
The *Ch*-induced fragmentation of actin filaments was suppressed in *adf4* and *ADF1-4Ri*. (**a**) Observation of GFP-ABD2-labeled AFs in *Ch*-infected Col-0, *adf4* and *ADF1-4Ri*. The merged images of GFP and bright field are shown. White dotted lines and asterisks indicate *Ch*-infected cells. White circles show appressoria (ap). Bar indicates 50 µm. (**b**) The percentages of *Ch*-infected cells with fragmented AFs. 9∼20 infected cells were observed per leaf, and 5 leaves for each line were analyzed in an experiment. The data show the average of three independent experiments. Comparison with Col-0 was performed by Student’s *t*-test. Asterisks indicate that there is a significant difference (* *P*<0.05).

### ACTIN mutants showed increased resistance to *Ch*

To further investigate the role of actin in the response to *Ch*, we examined *Ch* infection in *A. thaliana* actin mutants. Among the eight ACTIN genes encoded in the *A. thaliana* genome, *ACT2*, *ACT7*, and *ACT8* are expressed in vegetative tissues. ACT2 and ACT8 share high sequence identity and exhibit functional redundancy, while ACT7 has a distinctive role (Kandasamy *et al*., 2009). Mature leaves of *act2/8* and *act7* were inoculated with *Ch*, and lesion size was measured at 6 dpi. We found that lesion size was significantly reduced in both mutants relative to wild type (Fig. 4a and 4b). Compared with wild type, lesion size was reduced by 66% in *act2/8* and by 77% in *act7*. RT-qPCR analysis further confirmed suppression of *Ch* growth in these mutants, with *ChACT* expression reduced by 64% in *act2/8* and 84% in *act7* (Fig. 4c). We next examined *Ch* entry and secondary hyphae formation at 3 dpi. Both the *Ch* entry rate and the frequency of secondary hyphae formation were significantly reduced in *act2/8* and *act7* (Fig. 4d and 4e). The *Ch* entry rate was reduced by 64% in *act2/8* and by 50% in *act7* relative to wild type. In addition, the frequency of secondary hyphae formation was reduced by more than 50% in both mutants. Taken together, these results indicate that resistance to *Ch* is also enhanced in actin mutants.

**Figure 4.**
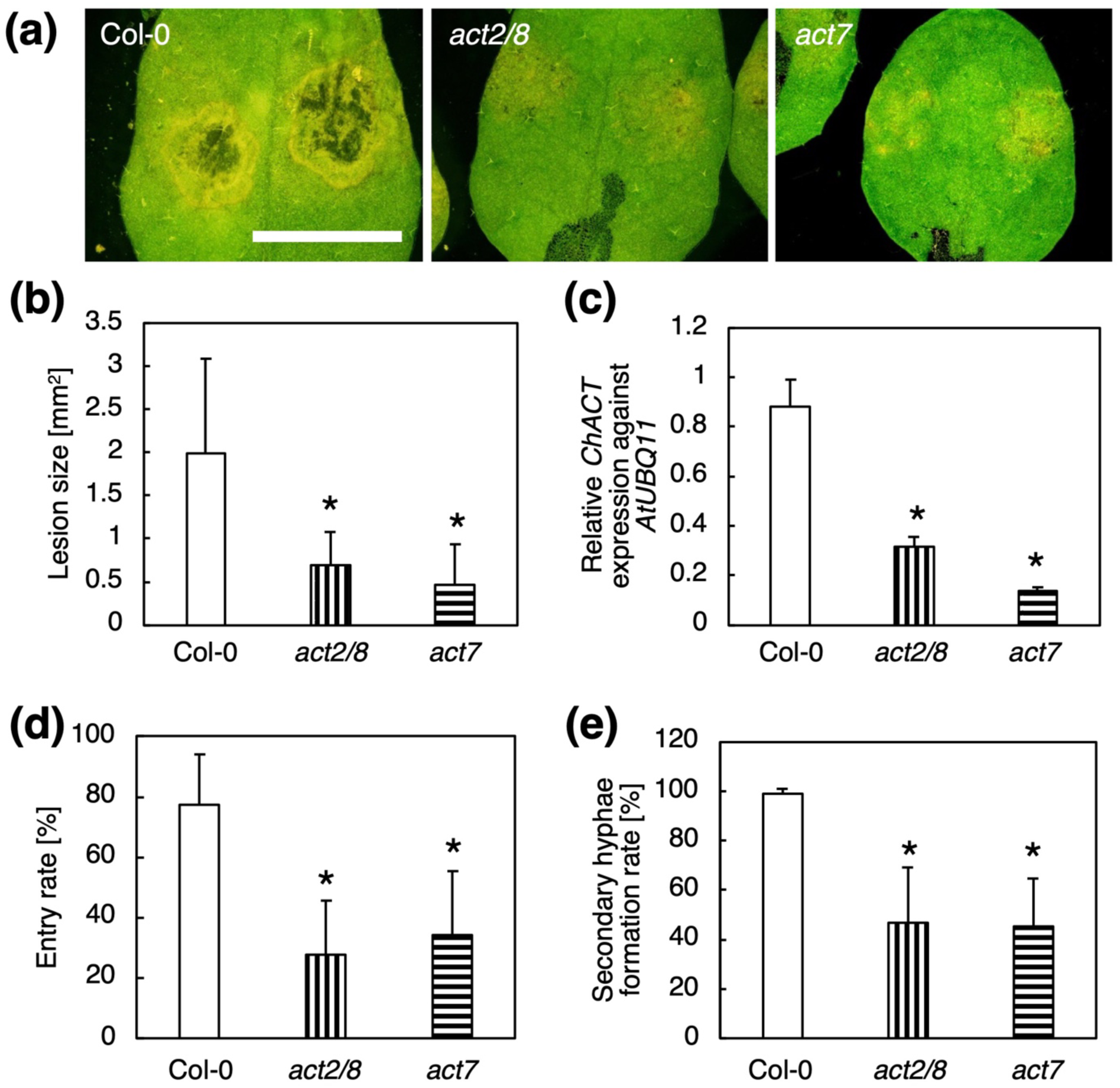
*A. thaliana act* mutants also showed an increased *Ch* resistance. (**a**) Col-0, *act2/8*, and *act7* mature leaves inoculated with *Ch* at 6 dpi. Bar indicates 5 mm. (**b**) The size of *Ch*-induced lesion at 6 dpi. The data showed average and standard deviation of 28 lesions for Col-0, *act2/8*, and *act7* each. (**c**) Quantification of *Ch* in planta by qRT-PCR. For (**b**) and (**c**), experiments were repeated three times with similar results. The representative results are shown. (**d**) *Ch* entry rate at 3 dpi. The percentage of appressoria with primary hyphae was counted for 20 appresoria per a leaf, and 16 leaves were used to calculate averaged percentages of entry. (**e**) The secondary hyphae formation rate at 3 dpi. The percentage of *Ch* forming secondary hyphae was determined among *Ch* forming primary hyphae. 20 *Ch* were counted per a leaf, and 18 leaves were used to calculate averaged percentages of secondary hyphae formation. For (**b**) to (**e**), comparison with Col-0 was performed by Student’s *t*-test. Asterisks indicate that there is a significant difference (\**P*<0.05).

### PEN2 functions in the increased *Ch* resistance of *adf4* and *ADF1-4Ri*

*A. thaliana* exhibits strong resistance to non-adapted *Colletotrichum* species, including *C. lagenarium*, *C. truncatum*, and *C. gloeosporioides*. Penetration attempts by these non-adapted pathogens are restricted by papilla formation, a dome-shaped cell wall structure generated through callose deposition at attempted penetration sites. This callose accumulation requires AFs, which become polarized beneath the appressoria of non-adapted *Colletotrichum* species (Shimada *et al*., 2006).

To determine whether the increased resistance to *Ch* entry observed in *adf4* and *ADF1-4Ri* is associated with enhanced callose deposition at *Ch* penetration sites, we performed aniline blue staining of *Ch*-infected leaves (Fig. S8a). Previous work demonstrated callose accumulation at attempted penetration sites of non-adapted *Colletotrichum* species at 1 dpi (Shimada *et al*., 2006). In Col-0, aniline blue-stained callose was associated with 5% of *Ch* appressoria at 1 dpi (Fig. S8b) and with 12% at 2 dpi (Fig. S8c). Both *adf4* and *ADF1-4Ri* exhibited levels of callose deposition comparable to those observed in Col-0 at both time points (Fig. S8a-c).

PEN2, an atypical myrosinase that hydrolyzes indole glucosinolates (Bednaerk *et al*., 2009), functions in penetration resistance to non-adapted *Colletotrichum* species (Hiruma *et al*., 2010; Irieda & Takano, 2021). PEN2 also contributes to penetration resistance against the *Ch* mutants *path-27* and *path-28*, which exhibit significantly reduced entry into host *A. thaliana* cells (Huser *et al*., 2009). To examine the involvement of PEN2 in the enhanced *Ch* resistance observed in *adf4* and *ADF1-4Ri*, we generated the double mutants *pen2-1 adf4* and *pen2-1*;*ADF1-4Ri* and inoculated them with *Ch* together with Col-0, *pen2-1*, *adf4*, and *ADF1-4Ri* controls. The lesion size of *pen2-1* was comparable to that of Col-0 (Fig. 5a-c). Introduction of the *pen2-1* mutation significantly increased lesion size in both *adf4* and *ADF1-4Ri*, and lesion sizes in the double mutants were comparable to those of Col-0 and *pen2-1* (Fig. 5a-c). We also analyzed *Ch* entry and secondary hyphae formation. As shown in Fig. 5d and 5e, introduction of the *pen2-1* mutation restored both the *Ch* entry rate and the secondary hyphae formation rate in *adf4* and *ADF1-4Ri* to levels comparable to those observed in wild type and *pen2-1*.

**Figure 5.**
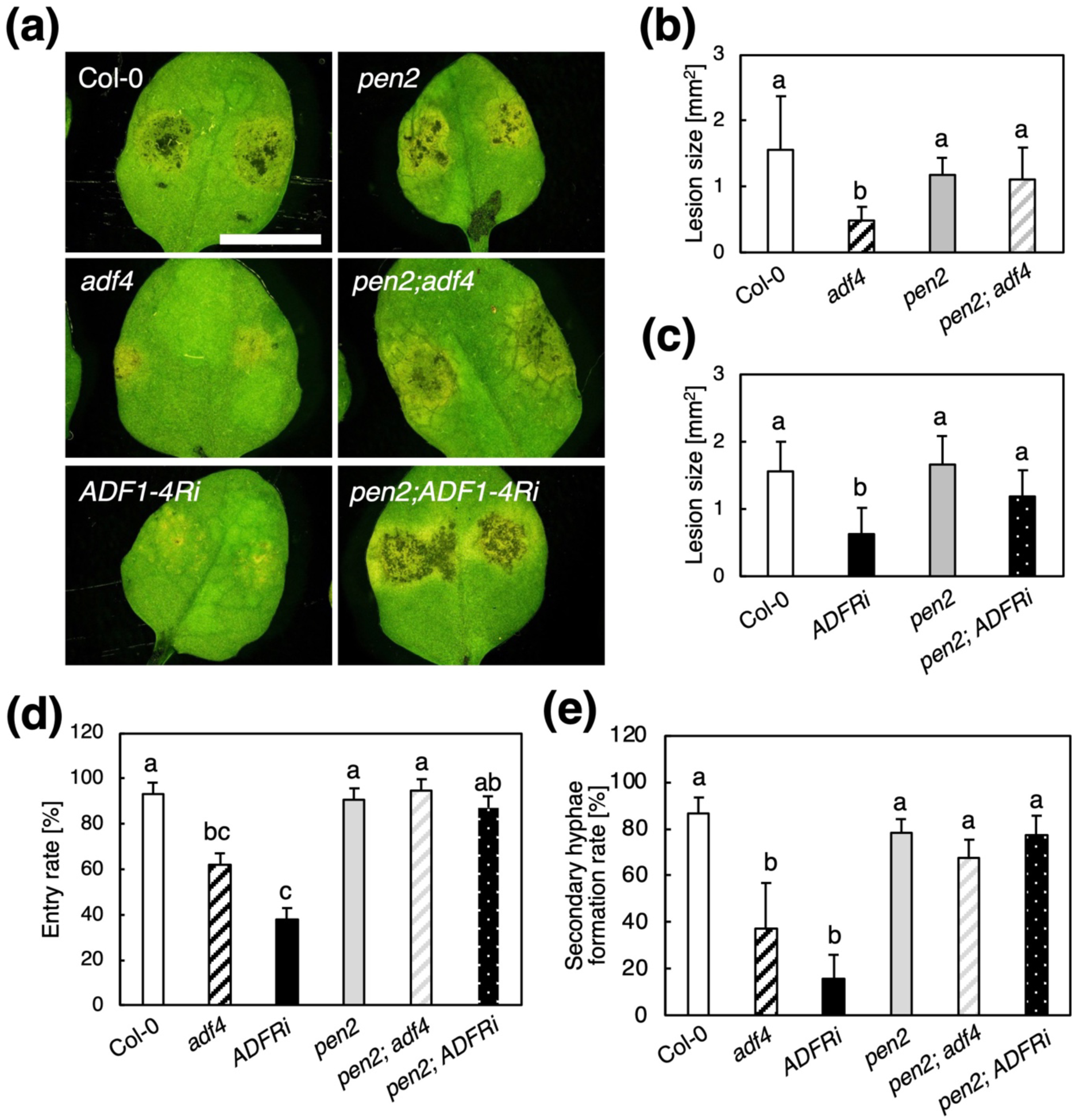
PEN2 plays a critical role in *Ch* resistance in *adf4* and *ADF1-4Ri*. (**a**) Col-0, *pen2*, *adf4*, *pen2*;*adf4*, *ADF1-4Ri*, and *pen2*;*ADF1-4Ri* mature leaves inoculated with *Ch* at 6 dpi. Bar indicates 5 mm. (**b**, **c**) The size of *Ch*-induced lesion at 6 dpi. The data showed average and standard deviation of 15 lesions for Col-0, *adf4*, *pen2,* and *pen2;adf4* each (**b**) and 16 lesions for Col-0, *ADF1-4Ri*, *pen2,* and *pen2;ADF1-4Ri* each (**c**). (**d**) *Ch* entry rate at 3 dpi. The percentage of appressoria with primary hyphae was counted for 20 appresoria per a leaf, and 7 leaves were used to calculate averaged percentages of invasion. (**e**) The secondary hyphae formation rate at 3 dpi. The percentage of *Ch* forming secondary hyphae was determined among *Ch* forming primary hyphae. 20 *Ch* were counted per a leaf, and 7 leaves were used to calculate averaged percentages of secondary hyphae formation. For (**b**) to (**e**), Comparisons among multiple groups were performed by ANOVA following the Turkey-Kramer test for each data set. The same letter indicates that there are no significant differences (*P*<0.05).

PEN2 accumulates at attempted penetration sites of non-adapted *C. gloeosporioides* (Hiruma *et al*., 2010). To determine whether PEN2 accumulation at *Ch* penetration sites is enhanced in *adf4* and *ADF1-4Ri*, we crossed PEN2-GFP-expressing Col-0 plants with *adf4* and *ADF1-4Ri*. However, PEN2-GFP fluorescence was markedly reduced in both *adf4* and *ADF1-4Ri*, likely due to transgene silencing, preventing direct analysis of PEN2 localization in these backgrounds.

Previous studies have shown that PEN2 localizes to both peroxisomes (Lipka *et al*., 2005) and mitochondria (Fuchs *et al*., 2016). In the interaction between *A. thaliana* and the non-adapted powdery mildew fungus *Blumeria graminis* f. sp. *hordei* (*Bgh*), mitochondria-associated PEN2 accumulates at sites of attempted fungal penetration (Fuchs *et al*., 2016). Because direct visualization of PEN2 localization using PEN2-GFP was not possible, we indirectly examined PEN2 localization in *adf4* and *ADF1-4Ri* by analyzing the distribution of peroxisomes and mitochondria. Because *Ch* entry began between 1 and 2 dpi under our experimental conditions, leaves collected at 30 hpi were used for these observations.

Observation of GFP-PTS1-labeled peroxisomes (Mano *et al*., 2002) in *Ch*-infected cells at 30 hpi revealed no obvious accumulation of peroxisomes at *Ch* penetration sites (Fig. S9). We therefore examined mitochondrial localization using MitoTracker staining. MitoTracker-labeled mitochondria were associated with many *Ch* appressoria at 30 hpi (Fig. 6a). Approximately 25% of *Ch* appressoria in Col-0 were associated with mitochondrial accumulation. This percentage was significantly increased in both *adf4* and *ADF1-4Ri* (Fig. 6b), with increases of 86% and 66%, respectively.

**Figure 6.**
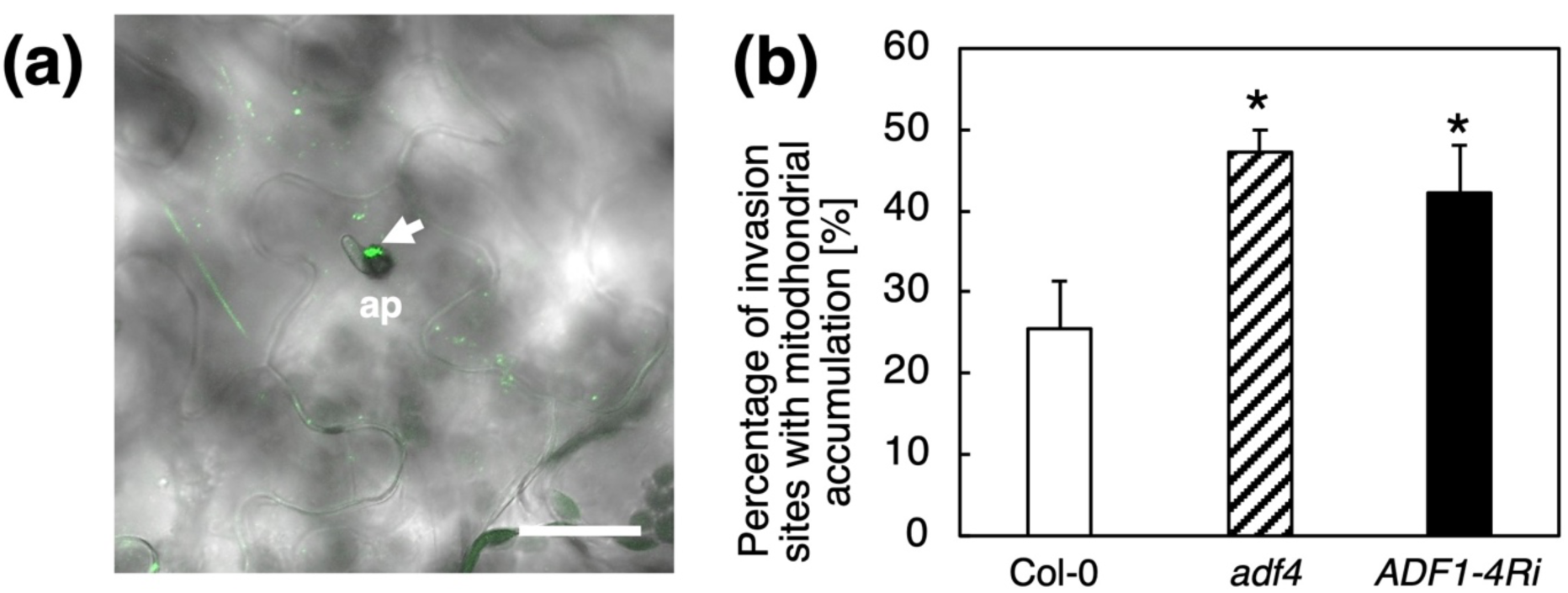
The mitochondrial accumulation at the *Ch* invasion site was increased in *adf4*. (**a**) Col-0 mature leaves at 30 hpi were inoculated with MitoTracker and observed using CLSM. The merged image of Alexa488 and bright field is shown. The arrow indicates the accumulation of mitochondria labeled by MitoTracker in an association with appressorium (ap). Bar indicates 50 µm. (b) The percentage of penetration sites with mitochondrial accumulation. 20∼60 conidiophores with appressoria were counted per sample in one experiment, and the averages of four independently performed experiments were shown. Comparison with Col-0 was performed by Student’s *t*-test. Asterisks indicate that there is a significant difference (\**P*<0.05).

Taken together, these results suggest that PEN2 accumulation at *Ch* penetration sites is enhanced in *adf4* and *ADF1-4Ri*, thereby suppressing *Ch* entry into host cells.

## DISCUSSION

In this study, we demonstrated that loss of subclass I ADFs in *A. thaliana* increases resistance to *Ch*. In *adf4* and *ADF1-4Ri*, this enhanced resistance was observed at both the *Ch* entry stage and the stage of secondary hyphae formation. *Ch*-induced AF fragmentation was significantly suppressed in both *adf4* and *ADF1-4Ri*, and, actin mutants also exhibited increased resistance to *Ch*. Introduction of the *pen2-1* mutation into *adf4* and *ADF1-4Ri* completely abolished the enhanced *Ch* resistance observed in these plants. Furthermore, we observed accumulation of mitochondria, to which PEN2 localizes, at *Ch* entry sites in both *adf4* and *ADF1-4Ri*, indicating that PEN2 contributes to resistance against *Ch* entry in these plants.

Based on these findings, we propose a model explaining how loss of subclass I ADFs enhances resistance to *Ch* (Fig. 7). As previously discussed (Shimada *et al*., 2026), *Ch* likely promotes host AF fragmentation through the secretion of effector(s). PEN2-associated mitochondria move throughout the cell along AFs. In wild-type *A. thaliana*, *Ch*-induced AF fragmentation interferes with the accumulation of PEN2-associated mitochondria at *Ch* entry sites, thereby suppressing PEN2-mediated penetration resistance and facilitating successful infection. In contrast, *Ch*-induced AF fragmentation is suppressed in *adf4* and *ADF1-4Ri*, allowing efficient delivery of PEN2-associated mitochondria to *Ch* entry sites. The resulting accumulation of PEN2 at these sites suppresses *Ch* invasion.

**Figure 7.**
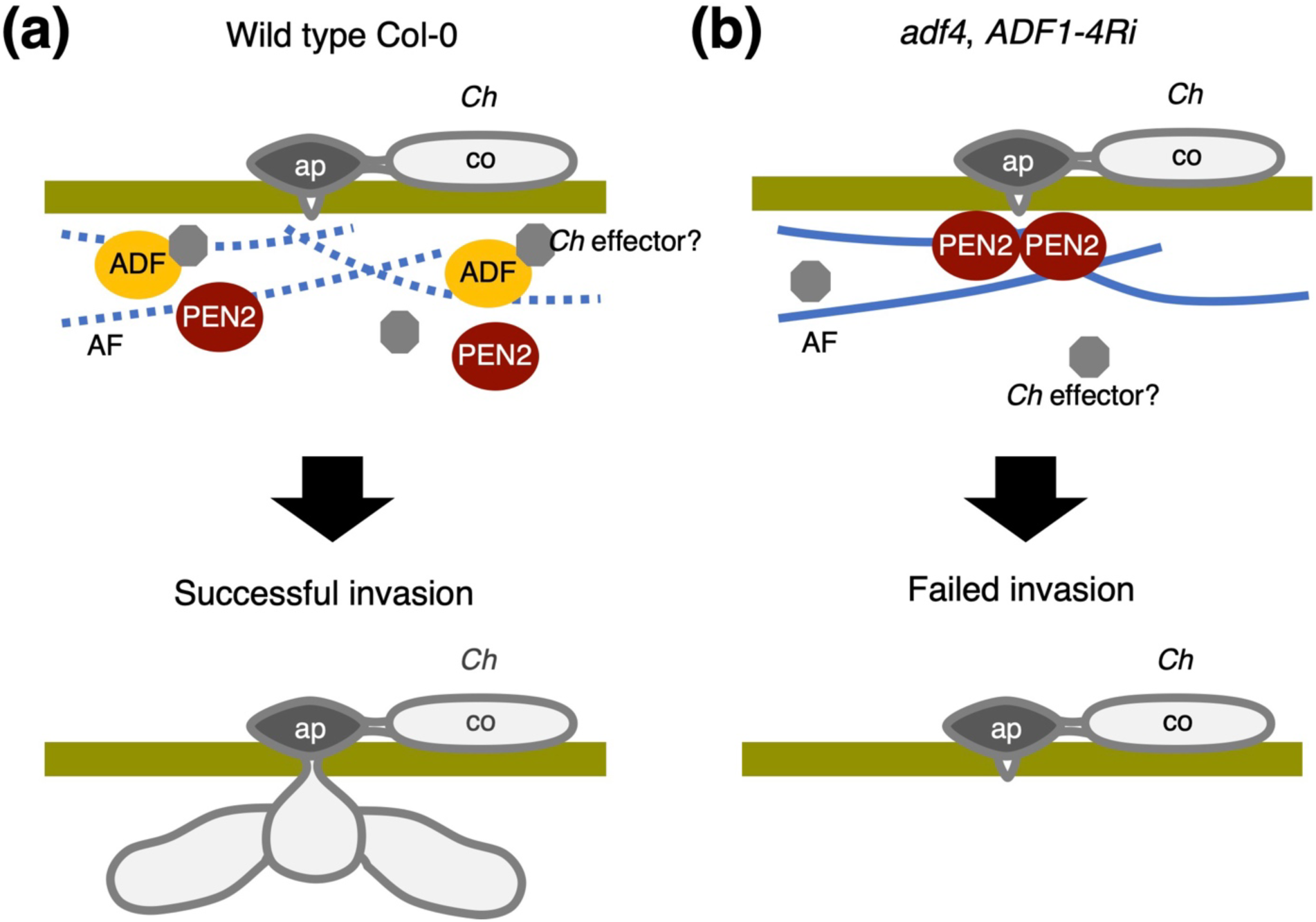
A working model for the mechanism of *Ch* resistance in *adf4* and *ADF1-4Ri*. (**a**) In wild type, *Ch* secretes effectors to modulate ADF activity to promote AF fragmentation. This AF fragmentation interferes accumulation of PEN2-associated mitochondria at *Ch* entry site, thereby suppressing PEN2-mediated penetration resistance and facilitating successful infection. (**b**) In *adf4* and *ADF1-4Ri*, *Ch*-induced AF fragmentation is suppressed, allowing efficient delivery of PEN2-associated mitochondria to *Ch* entry sites. The resulting accumulation of PEN2 at these sites suppresses *Ch* invasion. ap, appressorium; co, conidium.

This model assumes the existence of *Ch* effector(s) that directly or indirectly modulate host ADF activity to promote AF fragmentation. Several effectors that manipulate host AFs and actin-binding proteins (ABPs) have been identified in bacterial pathogens. For example, XopR from *Xanthomonas campestris* directly binds *A. thaliana* Formin 1 (AtFH1), a protein involved in AF nucleation, and promotes AF nucleation when present at low levels. XopR contains an intrinsically disordered region (IDR) and undergoes liquid-liquid phase separation. This IDR is required for cluster formation with AtFH1 and for stimulation of AF nucleation. In addition, XopR promotes AF bundling and inhibits ADF activity (Sun *et al*., 2021a). Another bacterial effector, HopW1 from *P. syringae*, directly binds host actin and disrupts AFs, thereby impairing AF-dependent membrane trafficking. Through suppression of host membrane trafficking, HopW1 enhances the pathogenicity of *P. syringae* (Kang *et al*., 2014). To date, no *Ch* effectors that target host AFs or ABPs have been identified. ADF4 localizes to both cytoplasmic AFs and the nucleus (Inada *et al*., 2016), yet previous analyses of *Ch* effector candidates (ChECs) did not identify any effectors that localize to plant AFs. EST-based analyses identified 102 ChECs predicted to function extracellularly, to be unique to the *Colletotrichum* genus, and to be specifically expressed during the biotrophic stage (Kleeman *et al*., 2012). Among these, 61 ChECs were selected based on high expression during the early stages of infection, including penetration and biotrophic primary hyphae formation, and their subcellular localization was examined by transient expression of GFP-tagged proteins in *Nicotiana benthamiana* epidermal cells (Robin *et al*., 2018). Of these 61 ChECs, 40 showed distribution throughout the cytoplasm and nucleus, 9 localized specifically to the nucleus, 6 accumulated in small punctate structures later identified as Golgi bodies or peroxisomes, and 3 were associated with microtubules (Robin *et al*., 2018). ChECs expressed at lower levels, which were excluded from the localization study (Robin *et al*., 2018), may be involved in modulation of host ADF activity. Future studies aimed at identifying *Ch* effectors that target host AFs and ADFs will contribute to a better understanding of *Ch* infection mechanisms and may facilitate the development of strategies to control *Ch* disease.

In this study, *act* mutants also showed increased *Ch* resistance similarly to *adf4* and *ADF1-4Ri*. This result may be explained by an alteration of AF architecture caused by loss of ACT isoform. Previously, a transient expression assay showed that AFs consist of either a single actin isoform (ACT2 or ACT7) or a mixture of both in *A. thaliana* protoplasts (Kijima *et al*., 2018). In *Nicotiana benthamiana* mesophyll cells, although both ACT2 and ACT7 are incorporated into the same filaments, they exist in distinct patch-like distributions. Furthermore, in the *N. benthamiana* epidermal cells, ACT2 localizes to fine filaments whereas ACT7 localizes to bundled thick filaments (Kijima *et al*., 2018). The same group has also been reported that different ACT isoforms exhibit distinct interactions with ABPs and differentially affect their activities (Kijima *et al*., 2016). Taking these findings into account, it is expected that the loss of a specific ACT isoform alters the physical composition of AFs, such as their isoform composition and bundling state. This alteration in AF composition may subsequently affect the binding and severing or depolymerizing activities of ADF, resulting in suppression of ADF activity and an increased resistance to *Ch*.

The *Ch*-resistant phenotype of *adf4* and *ADF1-4Ri* resembles that observed in several *Ch*-resistant *A. thaliana* accessions. Previous studies have shown that susceptibility to *Ch* varies considerably among *A. thaliana* accessions (Narusaka *et al*., 2004; Birker *et al*., 2009). Birker *et al*. (2009) reported that resistance in Ws-0, Gifu-2, Can-0, and Kondra was associated with suppression of *Ch* penetration and primary hyphae formation. These resistant accessions did not exhibit enhanced ROS accumulation in *Ch*-infected cells relative to susceptible accessions (Birker *et al*., 2009). Resistance in Ws-0, Gifu-2, Can-0, and Kondra was linked to the *RCH2* locus, which contains *RRS1* and *RPS4* (Birker *et al*., 2009; Narusaka *et al*., 2009). RRS1 and RPS4 are nucleotide-binding leucine-rich repeat (NB-LRR) proteins that directly recognize pathogen effectors. These proteins contribute not only to resistance against *Ch*, but also to resistance against *Ralstonia solanacearum* and *P. syringae* carrying AvrRps4 (Hinsch & Staskawicz 1996; Gassmann *et al*., 1999; Narusaka *et al*., 2009). Recognition of pathogen effectors by NB-LRR proteins is generally associated with activation of ETI and induction of cell death in infected cells (Jones & Dangl 2006). However, the mechanism underlying RCH2-associated resistance, which occurs without detectable cell death, remains unknown. It would therefore be of considerable interest to determine whether AF fragmentation and activation of the PEN2 pathway also occur in these *Ch*-resistant accessions, since this may provide insight into the mechanism underlying RCH2-associated *Ch* resistance.

In this study, loss of PEN2 suppressed not only resistance to *Ch* entry but also post-entry resistance, as the reduction in secondary hyphae formation observed in *adf4* and *ADF1-4Ri* was largely abolished by introduction of the *pen2-1* mutation (Fig. 5e). This dual role of PEN2 has been reported previously. Loss of PEN2 increases not only penetration by the non-adapted powdery mildew fungus *Bgh* but also subsequent hyphal elongation (Stein *et al*., 2006). *Ch* transitions to a necrotrophic phase through production of secondary hyphae from biotrophic primary hyphae (O’Connell *et al*., 2012), and PEN2-associated pathway is known to contribute to resistance against necrotrophic fungi. Loss of PEN2 and its downstream metabolites increases the susceptibility of *A. thaliana* to the necrotrophic fungus *Plectosphaerella cucumerina* (Sanchez-Vallet *et al*., 2010). PEN2 hydrolyzes 4-methoxyindol-3-ylmethylglucosinolate, a tryptophan-derived metabolite, generating indol-3-ylmethylglucosinolate, indol-3-ylmethylamine (I3A), and raphanusamic acid (RA) (Bednarek *et al*., 2009). *In vitro* treatment with either I3A or RA significantly suppresses germination and hyphal elongation of *P. cucumerina*, indicating that PEN2-derived metabolites possess antimicrobial activity (Sanchez-Vallet *et al*., 2010). PEN2 also contributes to resistance against the necrotrophic pathogen *Botrytis cinerea* (Buxdorf *et al*., 2013). We previously showed that expression of *PR1*, a marker gene for the salicylic acid defense pathway, is elevated in uninfected *ADF1-4Ri* plants (Inada *et al*., 2016), suggesting constitutive activation of basal defense. It is therefore possible that PEN2-mediated defense is also enhanced in *adf4* and *ADF1-4Ri*, contributing to suppression of post-entry *Ch* development.

We previously reported that *adf4* and *ADF1-4Ri* exhibit increased resistance to *Go*. In the present study, we identified several notable phenotypic differences between the enhanced resistance to *Ch* and that to *Go* in these plants. First, the elevated ROS accumulation observed in pathogen-infected *adf4* and *ADF1-4Ri* cells during *Go* infection was not detected during *Ch* infection (Fig. S4). Second, whereas ADF4-GFP-NES failed to complement the enhanced *Go* resistance phenotype of *adf4* (Inada *et al*., 2016), both ADF4-GFP and ADF4-GFP-NES complemented the increased *Ch* resistance phenotype of *adf4* (Fig. S5). Finally, expression of GFP-hTalin, which did not affect the enhanced *Go* resistance of *adf4* and *ADF1-4Ri* (Inada *et al*., 2016), significantly impaired the increased *Ch* resistance observed in these plants (Fig. S6). Collectively, these findings indicate that the mechanism underlying increased *Ch* resistance differs from that responsible for increased *Go* resistance in *adf4* and *ADF1-4Ri*.

As described in the Introduction, ADFs participate in a wide range of plant-pathogen interactions. In several cases, loss of ADF function results in altered AF organization at the cell surface. Clément et al. showed that inducible knockdown of *ADF2* increased AF bundling in root parenchyma cells, the tissue targeted by root-knot nematodes (Clément *et al*., 2009). Silencing of *TaADF3* altered AF orientation in wheat epidermal cells. In control plants, 78.4% of AFs were arranged longitudinally and 21.6% transversely, whereas the proportion of transversely oriented AFs increased to 39.8% in *TaADF3*-silenced plants (Tang *et al*., 2016). Suppression of *GhADF6* increased AF bundling and AF density in root cells and decreased the actin depolymerization rate in leaf epidermal cells of cotton plants (Sun *et al*., 2021b). Although these alterations in AF organization were thought to influence plant-pathogen interactions, the mechanism by which changes in AF architecture affect plant susceptibility remained unclear. In the present study, we provide evidence, for the first time, linking altered AF organization caused by loss of ADF function to changes in pathogen susceptibility.

Our study also demonstrates that ADFs can influence plant-pathogen interactions through multiple mechanisms. Previous work from both our group and Day’s group showed that ADFs contribute to responses against *Go* and *Pst* AvrPphB through regulation of gene expression. We previously demonstrated that the nuclear function of ADF4 is important for the response to *Go* (Inada *et al*., 2016). In addition, we reported that subclass I ADFs of *A. thaliana* participate in regulation of chromatin organization and gene expression (Matsumoto et al., 2023). Together, these findings suggest that altered gene expression in *adf4* and *ADF1-4Ri* contributes to enhanced immunity against *Go*. Consistent with this idea, expression of *RPS5*, which encodes the resistance protein recognizing the bacterial effector AvrPphB, was significantly reduced in *adf4*, resulting in increased susceptibility to *Pst* AvrPphB (Tian *et al*., 2009; Porter *et al*., 2012). More recently, the same group reported a novel role for ADF4 in gene regulation through interaction with transcription factors in the nucleus (Li *et al*., 2025). In contrast, our findings indicate that subclass I ADFs contribute to the response to *Ch* through their canonical function in regulation of cytoplasmic AFs. Thus, the diverse roles of ADFs in plant-pathogen interactions appear to be governed by their distinct functions in both the cytoplasm and the nucleus.

## ACKNOWLEDGEMENTs

We thank Prof. Brad Day at Michigan State University for his generous gift of *ADF1-4Ri* lines as well as GFP-ABD2 line We also thank Prof. Paul Schulze-Lefert at Max-Planck Institute and Prof. Yoshitaka Takano at Kyoto University for generous gifts of *pen2-1* and PEN2-GFP. GFP-PTS1 was kindly provided by Prof. Shoji Mano at NIBB. This work was supported by Grants-in-Aid for Scientific Research (16K07415, 20K06690 and 24K09518 for NI, and 26K02025 for TLS), the Osaka Prefecture University internal fund “RESPECT”, Osumi Frontier Science Foundation, the NOVARTIS Foundation (Japan) for the Promotion of Science, Nagase Science Technology Foundation, and Yamada Science Foundation (for NI), the NIBB Collaborative Research Program (25NIBB302 and 26NIBB303 for T.LS.), the Hamaguchi Foundation for the Advancement of Biochemistry, and Intensive Support Program for Young Promising Researchers from the New Energy and Industrial Technology Development Organization (NEDO) (24000770-0) (for TLS).

## AUTHOR CONTRIBUTIONS

NI planned and designed the research. NI, MO and SA performed experiments and data analyses. MU supervised the *Ch* infection analysis. TLS and TU supervised analysis of AF organization in *Ch*-infected leaves. NI wrote the manuscript with input from all authors. MO and SA contributed equally to this work.

## DATA AVAILABILITY

Raw data and/or original images will be made available to reviewers and the journal, if requested.

## REFERENCES

Bednarek P, Pislewska-Bednarek M, Svatoš A, Schneider B, Doubský J, Mansurova M, Humphry M, Consonni C, Panstruga R, Sanchez-Vallet A, et al. 2009. A glucosinolate metabolism pathway in living plant cells mediates broad-spectrum antifungal defense. Science 323:101–106.

Birker D, Heidrich K, Takahara H, Narusaka M, Deslandes L, Narusaka Y, Reymond M, Parker JE, O’Connell R 2009. A locus conferring resistance to *Colletotrichum higginsianum* is shared by four geographically distinct Arabidopsis accessions. Plant J 60:602–613.

Buxdorf K, Yaffe H, Barda O, Levy M 2013. The effects of glucosinolates and their breakdown products on necrotrophic fungi. PLOS ONE 8:370771.

Clément M, Ketelaar T, Rodiuc N, Banora MY, Smertenko A, Engler G, Abad P, Hussey PJ, de Almeida Engler J 2009. Actin-depolymerizing factor2-mediated actin dynamics are essential for root-knot nematode infection of *Arabidopsis*. Plant Cell 21: 2963–2979.

Collins NC, Thordal-Christensen H, Lipka V, Bau S, Kombrink E, Qiu J-L, Hückelhoven R, Stein M, Freialdenhoven A, Somervill SC, et al. 2003. SNARE-protein-mediated disease resistance at the plant cell wall. Nature 425:973–977.

Dean R, van Kan JAL, Pretorius ZA, Hammond-Kosack KE, di Pietro A, Spanu PD, Rudd JJ, Dickman M, Kahmann R, Ellis J, et al. 2012. The top 10 fungal pathogens in molecular plant pathology. Mol Plant Pathol 13:414–430.

Fu Y, Duan X, Tang C, Li X, Voegele RT, Wang X, Wei F, Kang Z. 2014. TaADF7, an actin-depolymerizing factor, contributes to wheat resistance against *Puccinia striiformis* f.sp. *tritici*. Plant J 78: 16–30.

Fuchs R, Kopischke M, Klapporodt C, Hause G, Meyer AJ, Schwarzländer M, Fricker MD, Lipka V 2016. Immobilized subpopulations of leaf epidermal mitochondria mediate PENETRATION2-dependent pathogen entry control in Arabidopsis. Plant Cell 28: 130–145.

Gassmann W, Hinsch ME, Staskawicz BJ 1999. The *Arabidopsis* RPS4 bacterial-resistance gene is a member of the TIR-NBS-LRR family of disease-resistance genes. Plant J 20:265–277.

Hinsch M, Staskawicz B 1996. Identificaiton of a new *Arabidopsis* disease resistance locus, RPS4, and cloning of the corresponding avirulence gene, *avrRps4*, from *Pseudomonas syringae* pv. *pisi*. Mol Plant Microb Interact 9: 55–61.

Hiruma K, Onozawa-Komori M, Takahashi F, Asakura M, Bednarek P, Okuno T, Schulze-Lefert P, Takano Y 2010. Entry mode-dependent function of an indole glucosinolate pathway in Arabidopsis for nonhost resistance against anthracnose pathogens. Plant Cell 22: 2429–2443.

Huser A, Takahara H, Schmalenbach W, O’Connell R 2009. Discovery of pathogenicity genes in the crucifer anthracnose fungus *Colletotrichum higginsianum* using random insertional mutagenesis. Mol Plant Microb Interact 22:143–156.

Inada N, Higaki T, Hasezawa S. 2016. Nuclear function of subclass I Actin Depolymerizing Factor contributes to susceptibility in Arabidopsis to an adapted powdery mildew fungus. Plant Physiol 170: 1420–1434.

Inada N. 2017. Plant Actin Depolymerizing Factor – Actin microfilament disassembly and more. J Plant Res 130: 227–238.

Irieda H, Takano Y 2021. Epidermal chloroplasts are defense-related motile organelles equipped with plant immune components. Nat Commun 12: 2739.

Jones JDG, Dangl JL 2006. The plant immune system. Nature 444:323–329.

Kandasamy MK, McKinney EC, Meagher RB 2009. A single vegetative actin isovariant overexpressed under the control of multiple regulatory sequences in sufficient for normal Arabidopsis development. Plant Cell 21: 701–718

Kang Y, Jelenska J, Cecchini NM, Li Y, Lee MW, Kover DR, Greenberg JT 2014. HopW1 from Pseudomonas syringae disrupts the actin cytoskeleton to promote virulence in Arabidopsis. PLOS Pathog 10: e1004232

Kijima ST, Hirose K, Kong S-G, Wada M, Uyeda TQP 2016. Distinct biochemical properties of *Arabidopsis thaliana* actin isoforms. Plant Cell Physiol 57: 46–46.

Kijima ST, Staiger CJ, Katoh K, Nagasaki A, Ito K, Uyeda TQP 2018. *Arabidopsis* vegetative actin isoforms, AtACT2 and AtACT7, generate distinct filament arrays in living plant cells. Sci Rep 8: 4381.

Kleeman J, Rincon-Rivera LJ, Takahara H, Neumann U, van Themaat EVL, van der Does HC, Hacquard S, Stüber K, Will I, Schmalenbach W et al. 2012. Sequential delivery of host-induced virulence effectors by appressoria and intracellular hyphae of the phytopathogen *Colletotrichum higginsianum*. PLOS Pathog 8:31002643.

Li P, Kelley B, Li Z, Procter B, Corrion A, Xie X, Sheick R, Lu Y-j, Nomoto M, Wei C-I, et al. 2025. Actin Depolymerization Factors (ADFs) moonlighting: nuclear immune regulation by interacting with WRKY transcription factors and shaping the transcriptome. Biorxiv doi: 10.1101/2025.04.29.651294

Lipka V, Dittgen J, Bednarek P, Bhat R, Wiermer M, Stein M, Landtag J, Brandt W, Rosahl S, Scheel D, et al. 2005. Pre- and postinvasion defences both contribute to nonhost resistance in *Arabidopsis*. Science 310:1180–1183.

Mano S, Nakamori C, Hayashi M, Kato A, Kondo M, Nishimura M. 2002. Distribution and characterization of peroxisomes in Arabidopsis by visualization with GFP: dynamic morphology and actin-dependent movement. Plant Cell Physiol 43: 331–341.

Matsumoto T, Higaki T, Takatsuka H, Kutsuna N, Ogata Y, Hasezawa S, Umeda M, Inada N. 2023. *Arabidopsis thaliana* subclass I ACTIN DEPOLYMERIZING FACTORs regulate nuclear structure and gene expression. Plant Cell Physiol 10: 1231–1242.

Mondal HA, Louis J, Archer L, Patel M, Nalam VJ, Sarowar S, Sivapalan V, Root DD, Shah J 2018. Arabidopsis *ACTIN-DEPOLYMERIZING FACTOR3* is required for controlling aphid feeding from the phloem. Plant Physiol 176: 879–890

Narusaka Y, Narusaka M, Park P, Kubo Y, Hirayama T, Seki M, Shiraishi T, Ishida J, Nakashima M, Enju A et al. 2004. *RCH1*, a locus in *Arabidopsis* that confers resistance to the hemibiotrophic fungal pathogen *Colletotrichum higginsianum*. Mol Plant Microb Interact 17:749–762.

Narusaka M, Shirasu K, Noutoshi Y., Kubo Y, Shiraishi T, Iwabuchi M, Narusaka Y 2009. RRS1 and RPS4 provide a dual Resistance-gene system against fungal and bacterial pathogens. Plant J 60: 218–226.

O’Connell R, Herbert C, Sreenivasaprasad S, Khatib M, Esquerré-Tugayé M-T, Dumas B 2004. A novel Arabidopsis-Colletotrichum pathosystem for the molecular dissection of plant-fungal interactions. Mol Plant Microb Interact 17:272–282

O’Connell RJ, Thon MR, Hacquard S, Amyotte SG, Kleemann J, Torres MF, Damm U, Buiate EA, Epstein L, Alkan N et al. 2012. Lifestyle transitions in plant pathogenic Colletotrichum fungi deciphered by genome and transcriptome analyses. Nat Genet 44:1060–1065

Porter K, Shimono M, Tian M, Day B 2012. Arabidopsis Actin-Depolymerizing Factor-4 links pathogen perception, defense activation and transcription to cytoskeletal dynamics. PLOS Pathogen 8: e1003006.

Robin GP, Kleeman J, Neumann U, Cabre L, Dallery J-F, Lapalu N, O’Connell RJ 2018. Subcellular localization screening of *Colletotrichum higginsianum* effector candidates identifies fungal proteins targeted to plant peroxisomes, Golgi bodies, and microtubules. Front Plant Sci 9:562.

Ruzicka DR, Kandasamy MK, McKinney EC, Burgos-Rivera B, Meagher RB 2007. The ancient subclasses of Arabidopsis ACTIN DEPOLYMERIZING FACTOR genes exhibit novel and differential expression. Plant J 52:460–472.

Saijo Y, Loo EP, Yasuda S 2018. Pattern recognition receptors and signaling in plant-microbe interactions. Plant J 93:592–613.

Sanchez-Vallet A, Ramos B, Bednarek P, López G, Pislewska-Bednarek M, Schulze-Lefert P, Molina A 2010. Tryptophan-derived secondary metabolites in *Arabidopsis thaliana* confer non-host resistance to necrotrophic *Plectrosphaerella cucumerina* fungi. Plant J 63:115–127.

Shimada C, Lipka V, O’Connell R, Okuno T, Shulze-Lefert P, Takano Y 2006. Nonhost resistance in *Arabidopsis*-*Colletotrichum* interactions acts at the cell periphery and requires actin filament function. MPMI 19: 270–279.

Shimada TL, Higaki T, Ebine K, Takano Y, Saijo Y, Ueda T 2026. Actin fragmentation induced by *Colletotrichum higginsianum* infection facilitates hyphal invasion in *Arabidopsis thaliana* leaves. Plant Cell Physiol doi: 10.1093/pcp/pcag038

Shimono M, Higaki T, Kaku H, Shibuya N, Hasezawa S, Day B. 2016. Quantitative evaluation of stomatal cytoskeletal patterns during the activation of immune signaling in *Arabidopsis thaliana*. PLoS One 11: e0159291.

Stein M, Dittgen J, Sángez-Rodríguez C, Hou B-H, Molina A, Schulze-Lefert P, Lipka V, Somerville S 2006. Arabidopsis PEN3/PDR8, an ATP binding cassette transporter, contributes to nonhost resistance to inappropriate pathogens that enter by direct penetration. Plant Cell 18:731–746.

Sun H, Zhu X, Li C, Ma Z, Han X, Luo Y, Yang L, Yu J, Miao Y 2021a. *Xanthomonas* effector XopR hijacks host actin cytoskeleton via complex coacervation. Nat Commun 12: 4064.

Sun Y, Zhong M, Li Y, Zhang R, Su L, Xia G, et al. 2021b. *GhADF6*-mediated actin reorganization is associated with defence against *Verticillium dahliae* infection in cotton. Mol Plant Pathol 22: 1656–1667.

Sun Y, Shi M, Wang D, Gong Y, Sha Q, Lv P, Yang J, Chu P, Guo S 2023. Research progress on the roles of actin-depolymerizing factor in plant stress responses. Front Plant Sci 14: 1278311.

Takahara H, Hacquard S, Kombrink A, Hughes B, Halder V, Robin GP, Hiruma K, Neumann U, Shinya T, Kombrink E, et al. 2016. *Colletotrichum higginsianum* extracellular LysM proteins play dual roles in appressorial function and suppression of chitin-triggered plant immunity. New Phytol 211:1323–1337.

Takemoto D, Jones DA, Hardham AR 2003. GFP-tagging of cell components reveals the dynamics of subcellular re-organization in response to infection of *Arabidopsis* by oomycete pathogens. Plant J 33: 775–792.

Tang C, Deng L, Chen S, Wang X, Kang Z 2016. TaADF3, an Actin-Depolymerizing Factor, negatively modulates wheat resistance against *Puccinia striiformis*. Front Plant Sci 6: 1214.

Tian M, Chaudhry F, Ruzicka DR, Meagher RB, Staiger CJ, Day B 2009. Arabidopsis Actin-Depolymerizing Factor AtADF4 mediates defense signal transduction triggered by the *Pseudomonas syringae* effector AvrPphB. Plant Physiol 150:815–824.

Yan Y, Yuan Q, Tang J, Huang J, Hsiang T, Wei Y, Zhang L 2018. *Colletotrichum higginsianum* as a model for understanding host-pathogen interactions. A review. Inter J Mol Sci 19:2142.

Zhang B, Hua Y, Wang J, Huo Y, Shimono M, Day B, Ma Q. 2017. *TaADF4*, an actin-depolymerizing factor from wheat, is required for resistance to the stripe rust pathogen *Puccinia striiformis* f. sp. *tritici*. Plant J 89: 1210–1224.

