## Supplementary materials for "*Arabidopsis thaliana* ACTIN DEPOLYMERIZING FACTORs are novel susceptibility factors for *Colletotrichum higginsianum*"

### SUPPLEMENTARY METHODS

#### DAB staining

For observation of ROS accumulation, *Ch*-inoculated leaves were incubated in 1% (w/v) DAB aqueous solution (pH 3.8) for 8 h in the light, then transferred to 70% ethanol and finally 100% ethanol before staining with Trypan blue as described in the maintext. Stained leaves were examined with a light microscope (Nikon ECLIPSE E600) equipped with 40x objective lens (Plant Fluor 40x N.A. 0.75, Nikon) and CCD camera (ORCA-ER, Hamamatsu photonics, Japan).

#### Callose staining

For observation of callose accumulation, *Ch*-inoculated leaves were fixed with 100% ethanol for 2 days at R.T. until the chlorophyll was completely removed. Fixed leaves were then incubated in 0.07 M phosphate buffer for 30 min. at R.T., then with 0.05% aniline blue solution [aniline blue (FUJI FILM, Osaka, Japan) was dissolved in 0.07 M phosphate buffer] for 1 hour at R.T..

Stained leaves were examined with a light microscope (Nikon ECLIPSE E600) equipped with 40x objective lens (Plant Fluor 40x N.A. 0.75, Nikon) and CCD camera (ORCA-ER, Hamamatsu photonics, Japan).

**Observation of lysosomes**

4 week old leaves of GFP-PTS1 plants were inoculated with *Ch* and observed with confocal laser scanning microscope (LSM700, Zeiss, Germany) equipped with 20x objective lens (Plan-Apochromat 20x DICII N.A. 0.8, Zeiss, Germany). GFP was excited with a 488 nm argon laser, and the emitted fluorescence was filtered with 505-600 nm bandpass filter.

**Table S1.** Primers used for RT-qPCR

| Gene | Sequence |
|---|---|
| ChACT F | 5'-CTCGTTATCGACAATGGTTC-3' |
| ChACT R | 5'-GAGTCCTTCTGGCCCATAC-3' |
| UBQ11 F | 5'-ACCAGCAGCGTCTCATCTTC-3' |
| UBQ11 R | 5'-TGTAGTCGGCCAAAGTACGTC-3' |

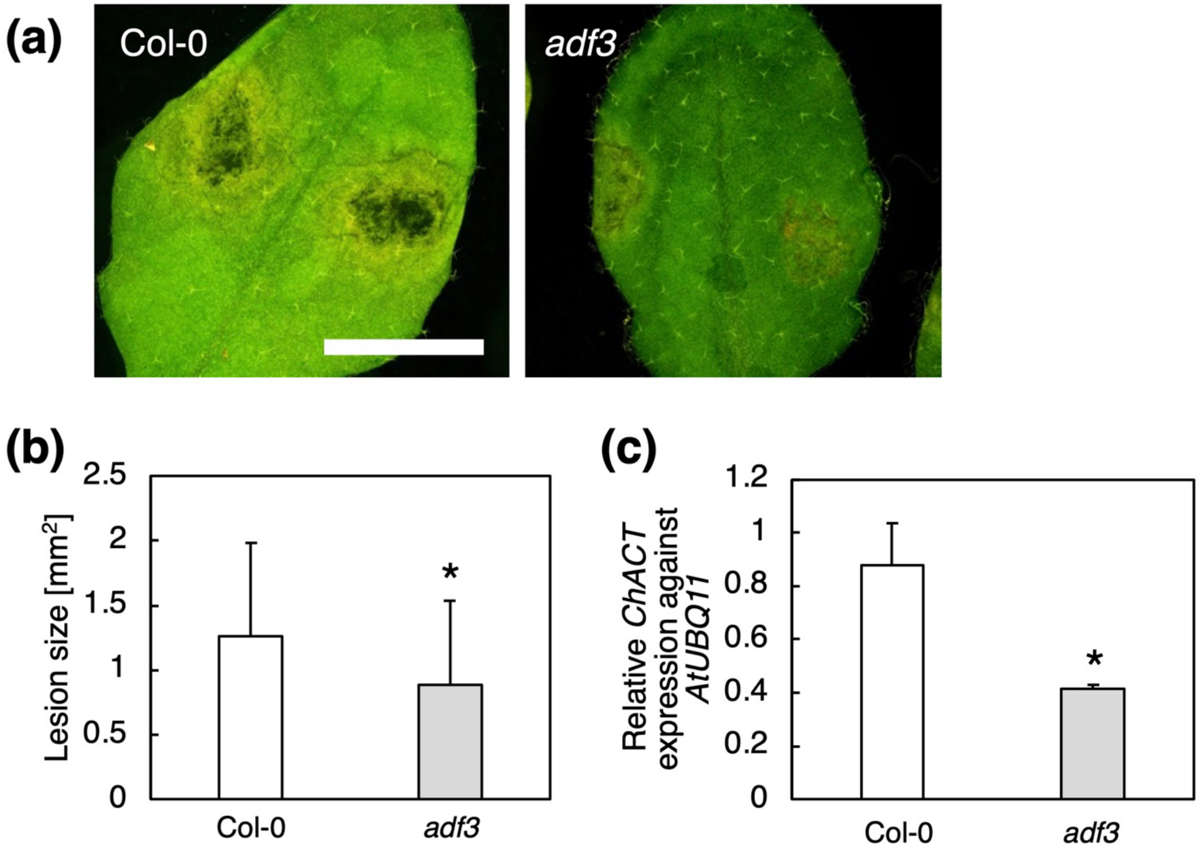


**Fig. S1** *Arabidopsis thaliana adf3* knockout mutant also showed an increased resistance against *C. higginsianum* (*Ch*). (**A**) Col-0 and *adf3* mature leaves inoculated with *Ch* at 6 days post inoculation (dpi). Bar indicates 5 mm. (**B**) The size of *Ch*-induced lesion at 6 dpi. The data showed average and standard deviation of 36 lesions for Col-0 and *adf3* each. (**C**) Quantification of *Ch* in planta by qRT-PCR. For (**B**) and (**C**), experiments were repeated three times with similar results. The representative results are shown. Comparison with Col-0 was performed by Student's *t*-test. Asterisks indicate that there is a significant difference (*$P<0.05$).

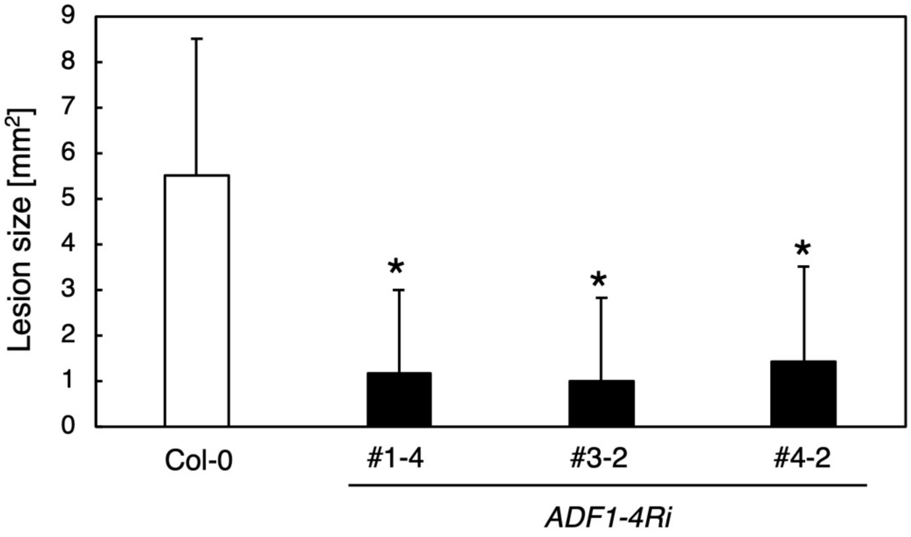


**Fig. S2** Multiple *ADF1-4Ri* lines showed an increased resistance against *Ch*. The size of *Ch*-induced lesion at 6 dpi. The data showed average and standard deviation of 20~30 lesions for Col-0 and three *ADF1-4Ri* lines. Comparison with Col-0 was performed by Student's *t*-test. Asterisks indicate that there is a significant difference (**P*<0.05).

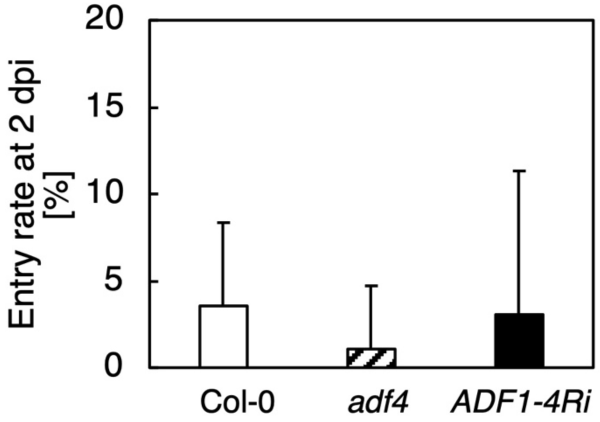


**Fig. S3** *Ch*-entry rate was unchanged in *adf4* and *ADF1-4Ri* compared to Col-0 at 2 dpi. The percentage of appressoria with primary hyphae was counted for 20 appresoria per a leaf, and 18 leaves were used to calculate averaged percentages of entry.

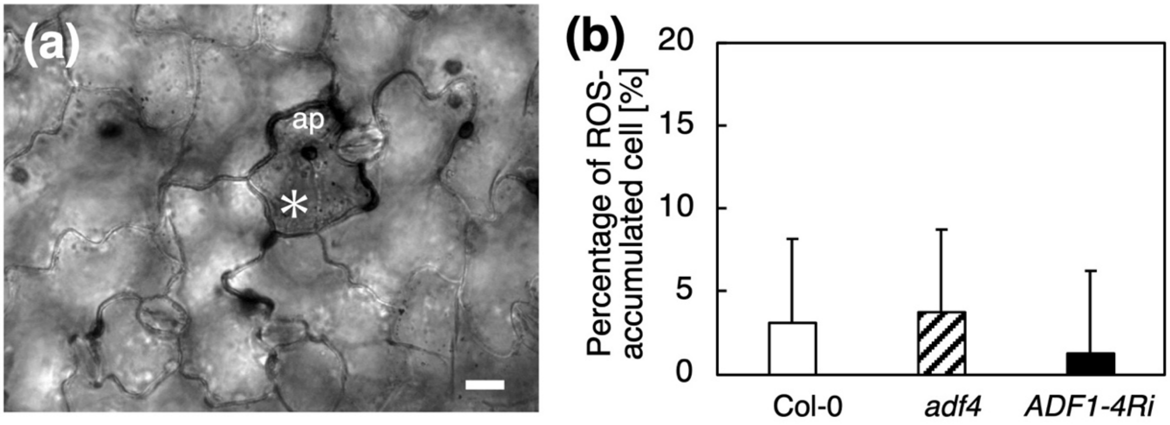


**Fig. S4** ROS-mediated resistance was unchanged in *adf4* and *ADF1-4Ri*. (**a**) *A. thaliana* mature leaves at 3 dpi was DAB-stained and observed under a microscope. Bar indicates 10 µm. ap, appressorium; *, *Ch*-infected cell. (**b**) The percentage of *Ch*-infected cells with ROS accumulation at 3 dpi. The percentage of ROS-accumulation was determined among Ch-infected cells. 20 *Ch*-infected cells were counted per a leaf, and 18 leaves were used to calculate averaged percentages. Comparison with Col-0 was performed by Student's *t*-test, and there were no significant differences ($P$<0.05).

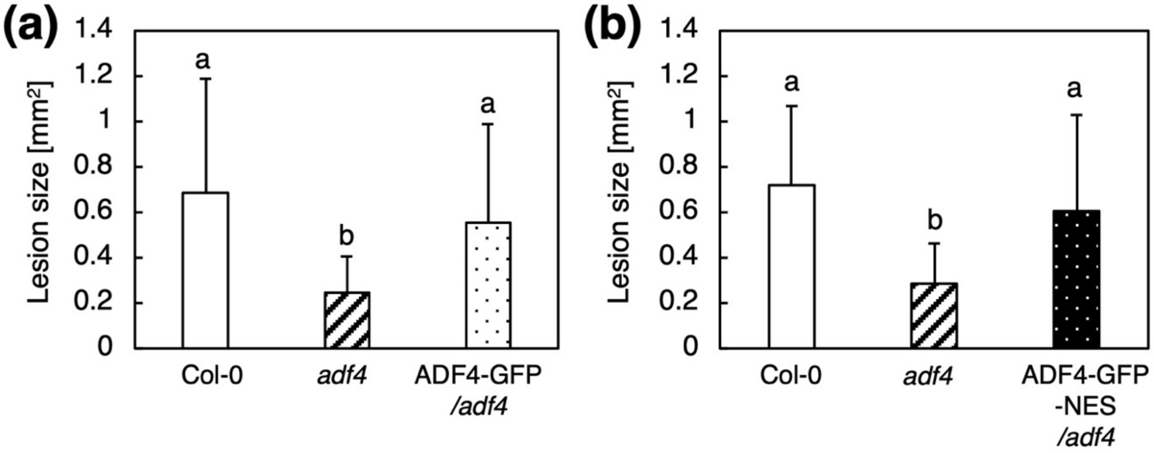


**Fig. S5** The nuclear localization of ADF4 was not important for response against *Ch*. (**a**) The size of *Ch*-induced lesion at 6 dpi. The data showed average and standard deviation of 30 lesions for Col-0, *adf4*, and ADF4-GFP/*adf4* each. (**b**) The size of *Ch*-induced lesion at 6 dpi. The data showed average and standard deviation of 30 lesions for Col-0, *adf4*, and ADF4-GFP-NES/*adf4* each. Comparisons among multiple groups were performed by ANOVA following the Turkey-Kramer test for each data set. The same letter indicates that there are no significant differences ($P$<0.05).

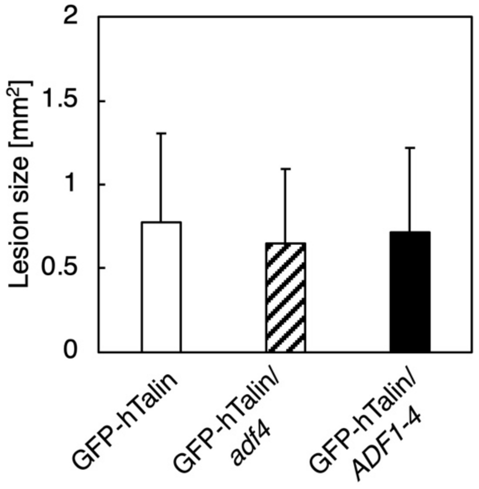


**Fig. S6** The expression of GFP-hTalin impaired *Ch* resistance in *adf4* and *ADF1-4Ri*. (**a**) The size of *Ch*-induced lesion at 6 dpi. The data showed average and standard deviation of 45 lesions for GFP-hTalin/Col-0, GFP-hTalin/*adf4*, and GFP-hTalin/*ADF1-4Ri* each. Comparison with Col-0 was performed by Student's *t*-test, and there were no significant differences ($P$<0.05).

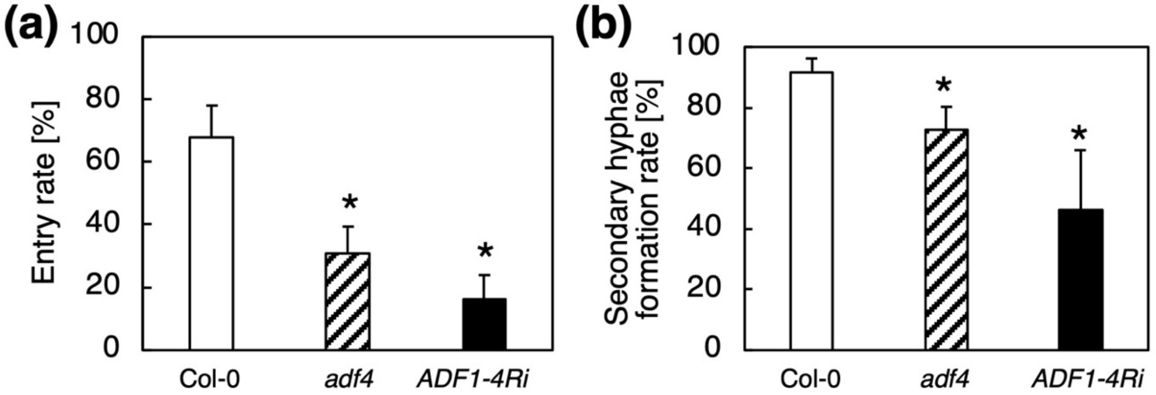


**Fig. S7** The increased *Ch* resistance was retained in cotyledons of *adf4* and *ADF1-4Ri*. (**a**) *Ch* entry rate at 3 dpi. The percentage of appressoria with primary hyphae was counted for 20 appresoria per a leaf, and 6 leaves were used to calculate averaged percentages of entry. (**b**) The secondary hyphae formation rate at 3 dpi. The percentage of *Ch* forming secondary hyphae was determined among *Ch* forming primary hyphae. 20 *Ch* were counted per a leaf, and 8 leaves were used to calculate averaged percentages of secondary hyphae formation. Comparison with Col-0 was performed by Student's *t*-test. Asterisks indicate that there is a significant difference (* $P$<0.05).

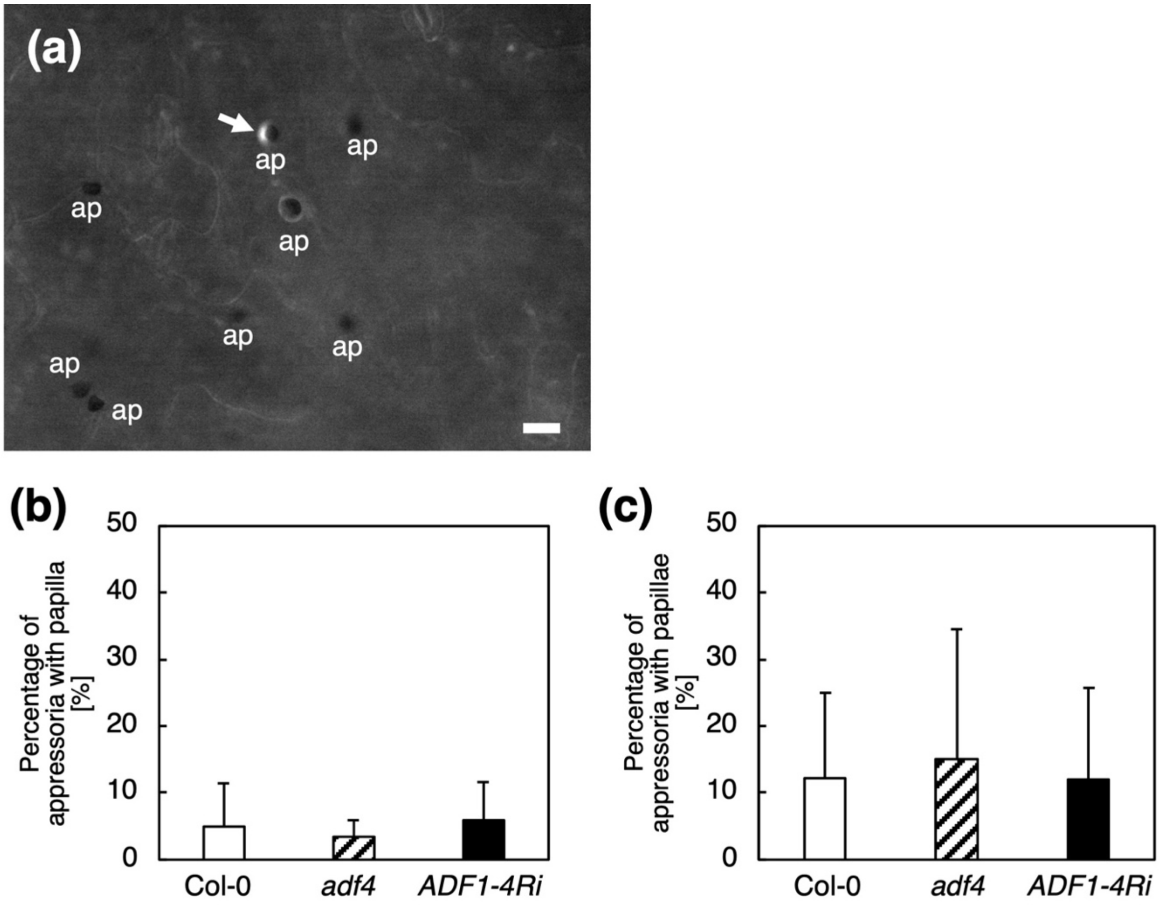


**Fig. S8** The callose accumulation at the *Ch* invasion site was not affected in *adf4* and *ADF1-4Ri*. (**a**) Leaves at 1-2 dpi were stained with aniline blue and observed under a fluorescence microscope. An arrow indicates the callose accumulation at the invasion sites. Bar indicates 10 µm. ap, appressorium. (b) The percentage of appressoria with callose accumulation at 2 dpi. The percentage of appressoria with callose accumulation was counted for 20 appresoria per a leaf, and 18 leaves were used to calculate averaged percentages. Comparison with Col-0 was performed by Student's *t*-test, and there was no significant difference ($P$<0.05).

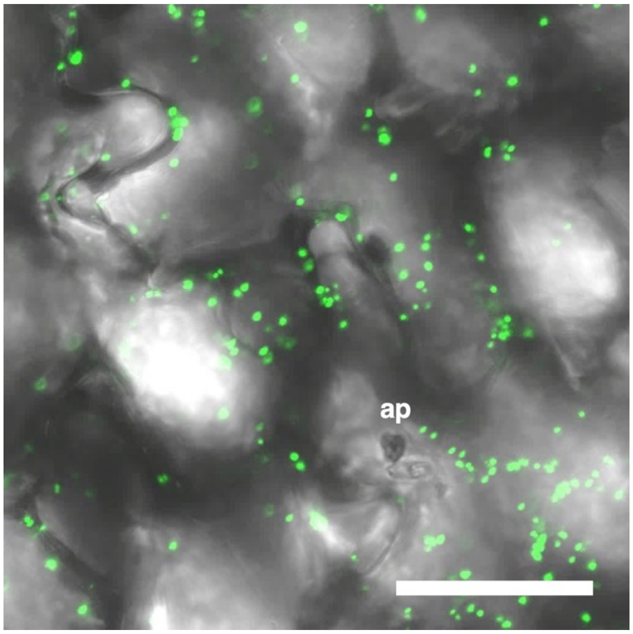


**Fig. S9** Localization of peroxisomes upon *Ch* invasion. Observation of GFP-PTS1-labeled peroxisomes in *Ch*-infected Col-0. The merged images of GFP and bright field are shown. ap, appressorium. Bar indicates 50 μm.
